# Cell division tracing combined with single-cell transcriptomics reveals new cell types and differentiation paths in the regenerating mouse lung

**DOI:** 10.1101/2023.01.18.524384

**Authors:** Leila R. Martins, Lina Sieverling, Michelle Michelhans, Chiara Schiller, Cihan Erkut, Sergio Triana, Stefan Fröhling, Lars Velten, Hanno Glimm, Claudia Scholl

**Author notes:** Corresponding authors, **Correspondence and requests for materials should be addressed to:** Claudia Scholl, Division of Applied Functional Genomics, DKFZ and NCT Heidelberg, Im Neuenheimer Feld 581, 69120 Heidelberg, Germany; or Leila Martins, Division of Applied Functional Genomics, DKFZ and NCT Heidelberg, Im Neuenheimer Feld 581, 69120 Heidelberg, Germany.

## Abstract

Understanding the molecular and cellular processes involved in lung epithelial regeneration may fuel the development of new therapeutic approaches for lung diseases. We combined new mouse models that allow diphtheria toxin (DTA)-mediated depletion of specific epithelial cell types and GFP-labeling of dividing cells with single-cell transcriptomics to characterize the regeneration of the distal lung. We uncovered new cell types, some of which likely represent epithelial precursors, propose goblet cells as progenitor cells, and provide evidence that adventitial fibroblasts act as supporting cells in epithelial regeneration. We also found that DTA-expressing cells can persist in the lung, express specific inflammatory factors, and resemble a previously undescribed population in the lungs of COVID-19 patients. Our study provides a comprehensive single-cell atlas of the distal lung that characterizes early transcriptional and cellular responses to defined epithelial injury, encompassing proliferation, differentiation, and cell-to-cell interactions.

## INTRODUCTION

Respiratory diseases are among the leading causes of death worldwide^1^. In the adult lung, epithelial cells possess a low steady-state turnover but can respond to lung injury with proliferation and differentiation to replace damaged cells^2^. Understanding the process of lung regeneration at the cellular and molecular level, including the identification of progenitor cells, is a prerequisite for the development of novel therapeutic strategies. Recent advances in single-cell transcriptomics of the human lung have enabled the identification of new cell types, increased the understanding of differentiation trajectories during lung development and regeneration, and allowed the comprehension of pathological processes from a cell type-specific point of view^3, 4, 5, 6, 7^.

Lung epithelial regeneration is commonly studied in mice, in which epithelial cells must first be injured to trigger repair. This is achieved by various means, including exposure to toxic gases^8, 9, 10^, detergents^11^, infectious agents^12^, chemicals^13,14^, and genetic approaches^13,15,16,17^. The advantage of genetic approaches is that they allow the depletion of specific cell types without interfering with the rest of the tissue.

The mouse trachea is lined by a pseudostratified epithelium containing ciliated cells, secretory goblet and club cells, basal cells, and rarer cell types such as tuft and neuroendocrine cells^2^. The morphology and cellular composition changes gradually from the trachea to the bronchioles, where the cuboid epithelium is composed mainly of ciliated and club cells. Rare bronchioalveolar stem cells (BASCs), co-expressing *Scgb1a1* and *Sftpc*, can be found in the bronchioalveolar duct junction. The alveoli are lined by cuboid alveolar type 2 cells (AT2) specialized in surfactant secretion, and flat alveolar type 1 cells (AT1) responsible for gas exchange^2^.

In the damaged lung, different epithelial cells can self-renew and differentiate, depending on the region and type of injury^18^. In the mouse trachea and main bronchi, basal cells are considered the major stem cell population^19^, while club cells are thought to act as progenitors for goblet^20^ and ciliated^9^ cells, and have been shown to regenerate the alveoli after injury with bleomycin or influenza^21^. BASCs have been reported to contribute to bronchioalveolar regeneration by their ability to differentiate into AT2, club, and ciliated cells^13^. Lastly, AT2 cells maintain the alveolar epithelium by self-renewing and differentiating into AT1 cells^15^.

Lung epithelial regeneration is also dependent on cues from the microenvironment^18^. For example, it has been shown that airway smooth muscle cells promote club cell repair by secreting Fgf10^22^, and a population of *Pdgfrα^+^* fibroblasts in the mouse trachea induced differentiation of basal cells through the production of Il6^23^. In the distal lung, *Axin2 Pdgfrα^+^* fibroblasts support AT2 cells through Il6, Bmp, and Fgf signaling modulation^24^. Using single-cell RNA sequencing (scRNA-seq), Tsukui *et al*. recently classified collagen-producing cells in the mouse and human lung as peribronchial fibroblasts, adventitial fibroblasts, alveolar fibroblasts, pericytes, and smooth muscle cells, but their role in epithelial regeneration is unclear^25^.

In spite of recent progress, there are still knowledge gaps on how the different lung cell types function during regeneration and whether there are any unknown cell types. In this study, we used scRNA-seq to analyze the consequences of epithelial cell type-specific depletion through inducible expression of diphtheria toxin subunit A (DTA) in the mouse distal lung, which has the potential to identify cell types or transition states that may be missed due to toxic effects after applying chemical or infectious agents. We combined this with a GFP-labeling approach to trace dividing cells *in vivo*, and applied a sorting strategy to enrich for rare cell types. This approach allowed us to identify new epithelial populations with potential progenitor cell properties, alternative differentiation trajectories, and adventitial fibroblasts as supporting cells during epithelial regeneration. We propose that club and ciliated cells participate in the damage-induced inflammatory response, and we show that viable DTA-expressing cells can persist in the lung and express a specific transcriptional profile that revealed a previously undescribed rare epithelial population in the lungs of COVID-19 patients.

## RESULTS

### *Scgb1a1^+^* cell depletion combined with *in vivo* tracing of dividing cells identifies early activated cell populations in the distal lung

To better understand which cells play an active role in lung epithelial regeneration, we generated a new mouse model that enables precise depletion of specific epithelial cell types and the isolation and detection of dividing cells. Specifically, we crossed Scgb1a1-CreER x Rosa26R-DTA mice^9^, in which tamoxifen administration induces DTA expression and subsequent apoptosis in *Scgb1a1*-expressing cells, with CycB1-GFP transgenic mice possessing GFP-labeled cells in the S and G2/M phases of the cell cycle^26^ (Fig. 1A). *Scgb1a1* (also known as *CC10*) is physiologically expressed by club cells and BASCs^9^. Scgb1a1-CreER x Rosa26R-DTA x CycB1-GFP (SRC) mice were injected with tamoxifen, and lungs were harvested before (day 0) and two and three days afterwards. Immunofluorescence (IF) analysis confirmed the progressive loss of club cells and revealed an increase in the number of GFP^+^ dividing cells over time (Fig. 1B). On day 2, the majority of GFP^+^ cells was located in the airway epithelium, consisting of club cells that were not yet damaged but replicated to repair the epithelium. On day 3, dividing GFP^+^ cells appeared in the airways, underlying connective tissue, and alveoli (Fig. 1B).

**Fig. 1:**
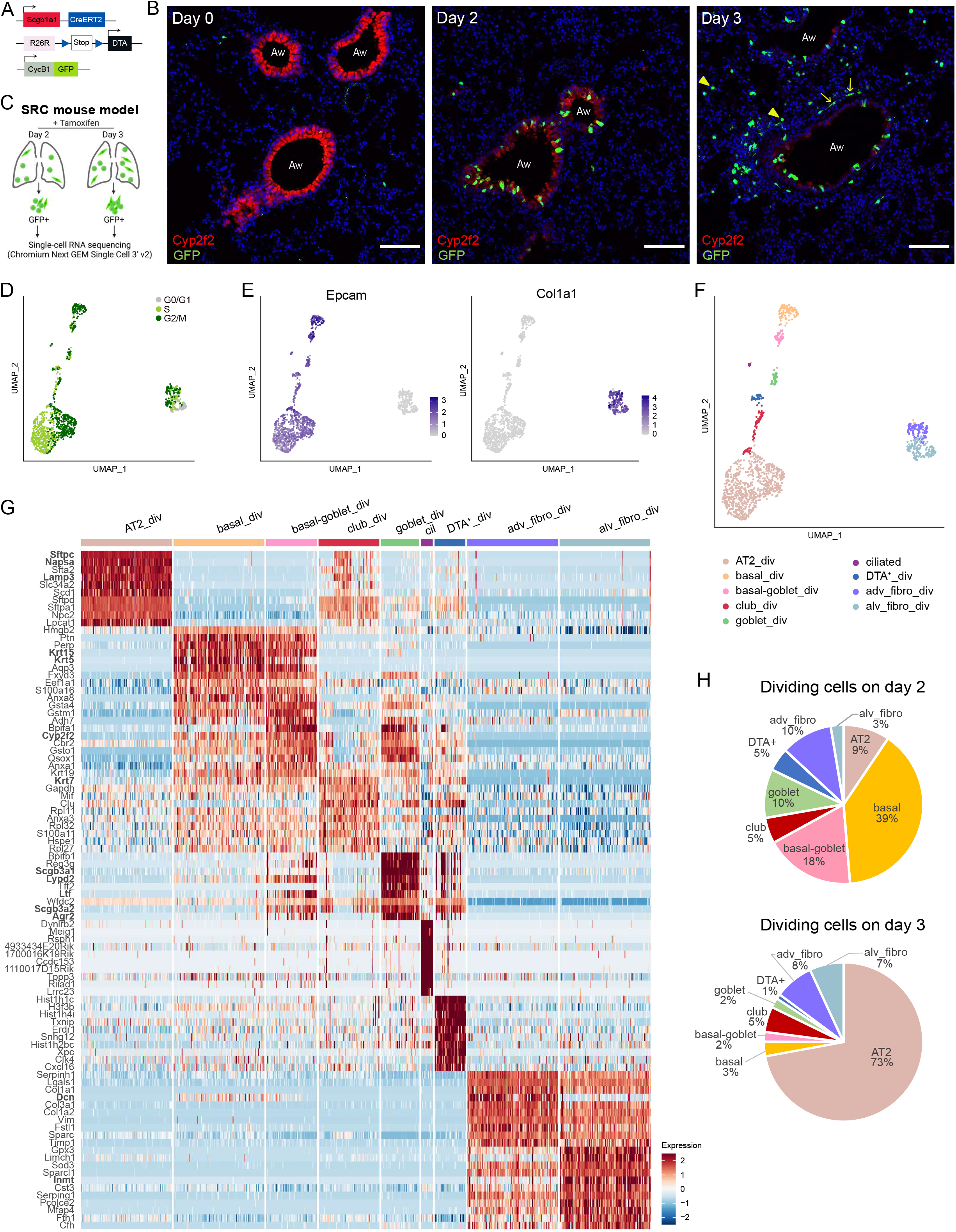
Characterization of dividing cells after targeted depletion of *Scgb1a1^+^* cells. **A.** Schematic of Scgb1a1-CreER, Rosa26R-DTA, and CycB1-GFP transgenes in SRC mice. **B**. IF staining of SRC mouse lungs with Cyp2f2 (red, club cells), GFP (green, dividing cells), and DAPI (blue, nuclei), before (day 0) and after 2 and 3 days of tamoxifen injection. GFP^+^ alveolar cells (arrow heads) and GFP^+^ spindle-like cells (arrows) can be observed on day 3. Aw: airway. Scale bar: 100 μm. Images are representative of at least three animals analyzed per group. **C.** Outline of cells used for scRNA-seq. Created with BioRender.com. **D-F.** UMAP embedding of scRNA-seq data from GFP^+^ cells sorted from lungs of SRC mice two days (n=3 mice) and three days (n=2 mice) after tamoxifen injection. **D.** Cell cycle phase distribution. **E.** Normalized expression of *Epcam* (epithelial cells) and *Col1a1* (mesenchymal cells). **F.** Cell type assignment. **G.** Heatmap of the top ten upregulated genes across cell populations ranked by power. Scaling of expression was done after downsampling to 100 cells per cell type. DEGs mentioned in the text are in bold. **H.** Percentage of dividing cell types in the distal lung at day 2 and 3 after tamoxifen injection.

To further characterize the heterogeneous population of proliferating cells that may contain stem or progenitor cells, we performed scRNA-seq of GFP^+^ cells isolated by fluorescent-activated cell sorting (FACS) from the distal lung of SRC mice at day 2 and 3 after tamoxifen injection (Fig. 1C). After excluding endothelial and hematopoietic cells, GFP^+^ cells represented 0.034% and 0.16% of total cells on day 2 and 3, respectively (Fig. S1A). *In silico* cell cycle analysis confirmed that the majority of sorted GFP^+^ cells were dividing, with 42% in G2/M and 44% in S phase (Fig. 1D), including both epithelial (*Epcam^+^*) and mesenchymal (*Col1a1^+^*) cells (Fig. 1E). Proliferating mesenchymal cells were identified as adventitial (Dcn^+^) and alveolar fibroblasts (Inmt^+^) (Fig. 1F and G). Five of the seven epithelial cell clusters could be allocated to distinct cell types based on the expression of known marker genes: AT2, basal, club, ciliated, and goblet cells (Fig. 1F and G, Supplementary Table 1). IF confirmed the transcriptomic results identifying GFP^+^ club, goblet, basal, and AT2 cells, as well as adventitial and alveolar fibroblasts (Fig. S1B). The small population of non-dividing ciliated cells (Fig. 1F) was isolated during FACS due to high autofluorescence. Basal cells represented 39% of dividing cells on day 2 (Fig. 1H), while AT2 cells accounted for the majority (73 %) of dividing cells on day 3 (Fig. 1H). This was unexpected, since AT2 cells are only known to function as progenitors for AT2 and AT1 cells upon alveoli injury ^2, 19^.

Among the differentially expressed genes (DEGs) in dividing club (club_div) cells compared to all other dividing cell types was the club cell marker *Krt7* (Fig. 1G)^27^, but other canonical club cell markers such as *Scgb3a2* or *Cyp2f2* were not consistently expressed (Fig. S1C). In fact, lower expression of these genes was associated with higher expression of AT2 marker genes including *Sftpc, Napsa*, and *Lamp3*, suggesting priming of club_div cells for differentiation into AT2 cells (Fig. 1G, Fig. S1C). Club_div cells showed no similarity to reported club cell progenitor populations including lineage-negative epithelial progenitors (*H2-K1^+^, Sox2^+^ Itgb4^+^*), variant club cells (Scgb3a2^high^, *Cyp2f2^-^, Upk3a^+^*), or hillock club cells (*Krt13^+^, Krt4^+^*) (Fig. S1C)^28,29,30^. Moreover, although 37% of club_div cells at day 3 co-expressed *Sftpc* and *Scgb1a1*, a characteristic feature of BASCs, they did not consistently express additional BASC marker genes described previously (Fig. S1D)^13,31^. This could suggest that either most BASCs were destroyed by DTA, or that they were not proliferating at the time points analyzed.

We found that goblet cells (*Agr2^+^, Ltf^+^, Scgb3a1^+^, Lypd2^+^*) which also expressed *Scgb1a1*, were dividing in the injured lung (Fig. 1F and G, Fig. S1F and Supplementary Table 1). This was surprising, since goblet cells have never been shown to be capable of self-renewal^19, 32^. Additionally, an undescribed cell type expressing both basal (*Krt5, Krt15*) and goblet (*Agr2, Lypd2, Ltf*) cell markers was found among the dividing cells, which we named basal-goblet_div, suggesting the existence of an intermediate progenitor population (Fig. 1F and G).

Finally, the remaining seventh cluster of dividing epithelial cells expressed DTA, indicating the presence of lung cells that survived the expression of this highly potent toxin (Fig. S1G). These DTA^+^ cells clustered separately from club_div and goblet_div cells and had a distinct gene signature, although they expressed secretory cell genes such as *Scgb3a2* (Fig. 1F and G, Supplementary Table 1).

In summary, *Scgb1a1*^+^ cell loss induces widespread cellular activation with immediate proliferation of several populations in the lung rather than triggering a specific cell type. Among the dividing cells were goblet cells and novel basal-goblet and AT2-primed club cell populations, which can represent new progenitors in the distal lung.

### Adventitial fibroblasts support lung epithelial regeneration

Several mesenchymal cell types are known to support epithelial regeneration upon lung injury^22, 23, 24^. Conversely, mesenchymal cells can also give rise to aberrant populations following lung injury, such as *Axin2^+^ Pdgfrα^-^* myogenic progenitors^24^ and *Cthrc1^+^* fibroblasts^25^, which contribute to fibrosis. To analyze the behavior of mesenchymal cells during depletion of *Scgb1a1^+^* cells, we performed scRNA-seq of dividing and non-dividing mesenchymal cells from the distal lung of SRC mice (Fig. 2A).

**Fig. 2:**
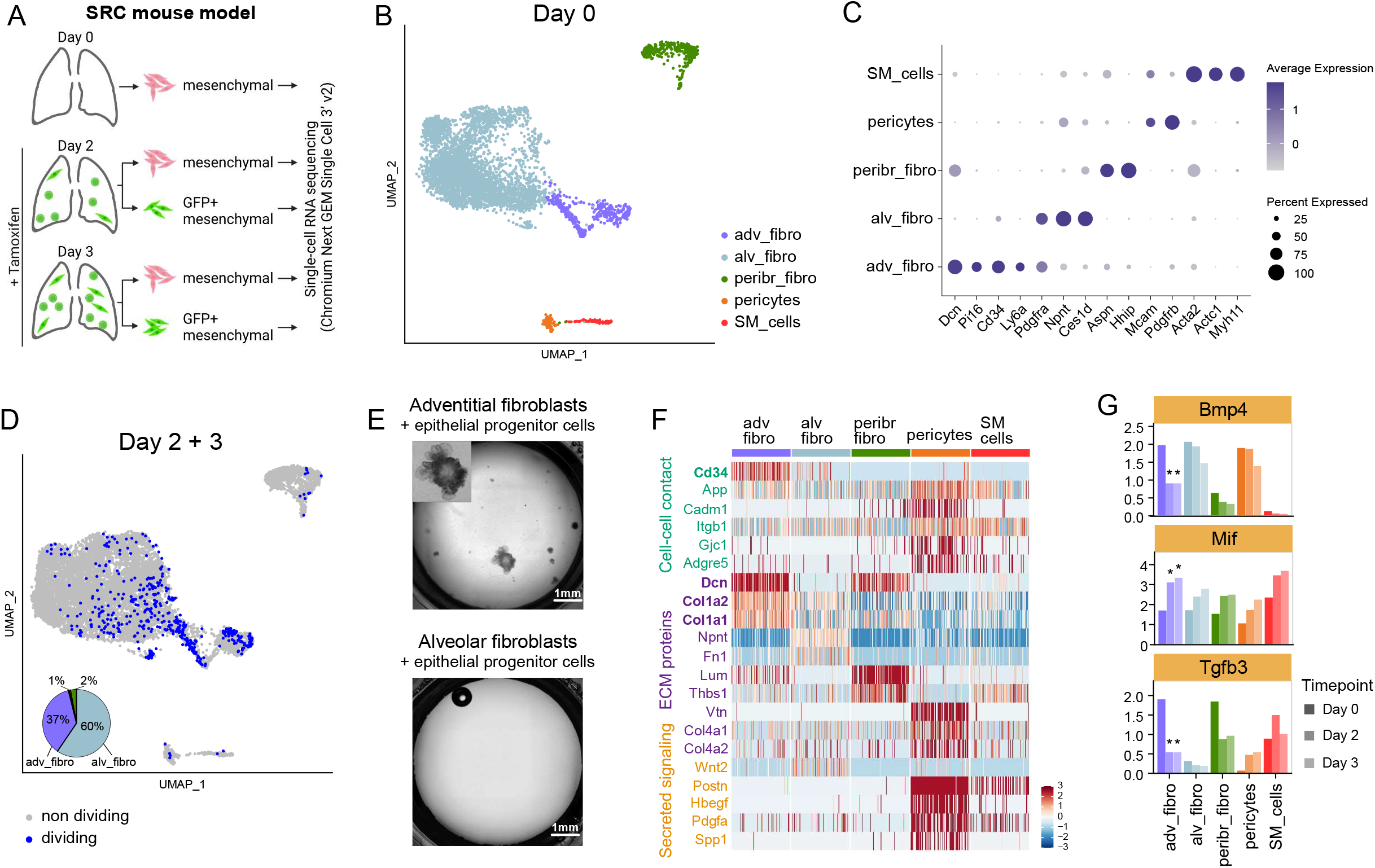
Characterization of mesenchymal cell activation after epithelial injury in SRC mice. **A.** Outline of cells used for scRNA-seq. Created with BioRender.com. **B.** UMAP embedding showing mesenchymal cell types sorted from uninjured SRC mouse lungs (n=2 mice). **C.** Dot plot showing the average scaled expression and percentage of cells expressing cell-type specific marker genes across the mesenchymal cell populations in uninjured lungs. **D.** UMAP embedding showing the distribution of dividing mesenchymal cells from GFP^+^ sorted cells on day 2 (n=12 mice) and day 3 (n=4 mice), and non-dividing total cells on day 2 (n=3 mice) and day 3 (n=2 mice). The pie chart depicts the percentage of the mesenchymal cell types among the diving cells (light blue: alveolar fibroblasts, purple: adventitial fibroblasts, green: peribronchial fibroblasts, orange: pericytes). **E.** Lung epithelial organoid cultures with adventitial fibroblasts or alveolar fibroblasts as supporting cells, taken after three weeks of culture. The inset in the upper image shows the morphology of one organoid magnified twice as much as in the main figure. Images are representative of two independent experiments. **F.** Heatmap of differentially expressed ligands among the different mesenchymal cell types on day 0. Upregulated genes in adventitial fibroblasts are bold. **G.** Average normalized expression of Bmp4, Tgfb3, and Mif on day 0, 2, and 3 in all mesenchymal populations. Asterisks denote differential expression on day 2 or 3 (FC >=1.5, p_adj < =0.05). adv_fibro: adventitial fibroblasts, alv_fibro: alveolar fibroblasts, peribr_fibro: peribronchial fibroblasts, SM_cells: smooth muscle cells.

In the uninjured lung, we identified adventitial fibroblasts, alveolar fibroblasts, peribronchial fibroblasts, pericytes, and smooth muscle cells (Fig. 2B and C, Supplementary Table 2). The types of mesenchymal cells remained unchanged upon injury (Fig. S2A), and none of the postinjury mesenchymal cell populations showed increased fibrosis-associated genes, suggesting that this mouse model does not induce lung fibrosis (Fig. S2B)^25, 33^. Analysis of the GFP^+^ cells confirmed our previous observation (Fig. 1E) that alveolar and adventitial fibroblasts are the major proliferating mesenchymal cell types after *Scgb1a1^+^* cell depletion, accounting for 60% and 37% of the total mesenchymal dividing cells, respectively (Fig. 2D). Since alveolar fibroblasts were approximately eight times more abundant than adventitial fibroblasts, the high proportion of adventitial fibroblasts in the total dividing cells suggests a higher activation and a possible functional role of these cells. To test this, we performed organoid cultures with epithelial progenitor cells (EPCAM^high^ CD24^dim^)^34^ in co-culture with adventitial fibroblasts (CD34^+^ SCA-1^+^) or alveolar fibroblasts (PDGFRA^+^ NPNT^+^) (Fig. S2C). After three to four weeks of culture, alveolar and bronchoalveolar organoids were formed with adventitial fibroblasts as supporting cells, but not with alveolar fibroblasts (Fig. 2E), reinforcing our hypothesis.

To learn more about the supporting role of adventitial fibroblasts, we examined the expression of genes encoding cell-cell contact proteins, extracellular matrix proteins and secreted ligands from the CellChat ligand-receptor database^35^. Adventitial fibroblasts differentially expressed *Cd34, Dcn, Col1a2*, and *Col1a1* in uninjured lungs (Fig. 2F), which was confirmed on protein level by mass spectrometry for CD34 and DCN, while COL1A1 was also high in alveolar fibroblasts (Fig. S2D). After injury, adventitial fibroblasts significantly downregulated mRNAs of the secreted ligands *Bmp4* and *Tgfb3* (Fig. 2G). These signaling proteins are known regulators of the lung epithelium. For example, inhibition of BMP signaling induces proliferation of lung epithelial cells *in vitro* and *in vivo*^36^ and TGF-β has a cytostatic effect on lung epithelial cells^37^. Adventitial fibroblasts also significantly upregulated several pro-inflammatory factors after *Scgb1a1^+^* cell depletion (Fig. 2G, Fig. S2E), corroborating the communication between mesenchymal and immune cells in the lung^38^. Among those was Mif (Fig. 2G), a pro-inflammatory factor that was reported to stimulate AT2 cell proliferation *in vitro*^39, 40^.

In summary, our data indicate that adventitial fibroblasts support epithelial repair as they proliferate extensively, produce pro-inflammatory factors, downregulate secreted ligands known to be cytostatic on lung epithelial cells, and support epithelial organoid growth in culture.

### Transcriptional changes after depletion of *Scgb1a1*\* cells suggest a role for club and ciliated cells in immune activation

A deeper understanding of the molecular changes in epithelial cells during lung regeneration can aid the development of therapies for lung diseases. We therefore characterized the cellular and transcriptional responses by scRNA-seq in dividing and non-dividing epithelial cells, sorted as Epcam^+^ cells from distal lungs of SRC mice without treatment and two, three, and four days after double tamoxifen administration (Fig. 3A). During sorting, AT2 cell numbers were reduced and GFP^+^ cells were enriched (Fig. S3A). The steady post-injury increase of GFP+ cells was confirmed by FACS and IF (Fig. S3A and D).

**Fig. 3:**
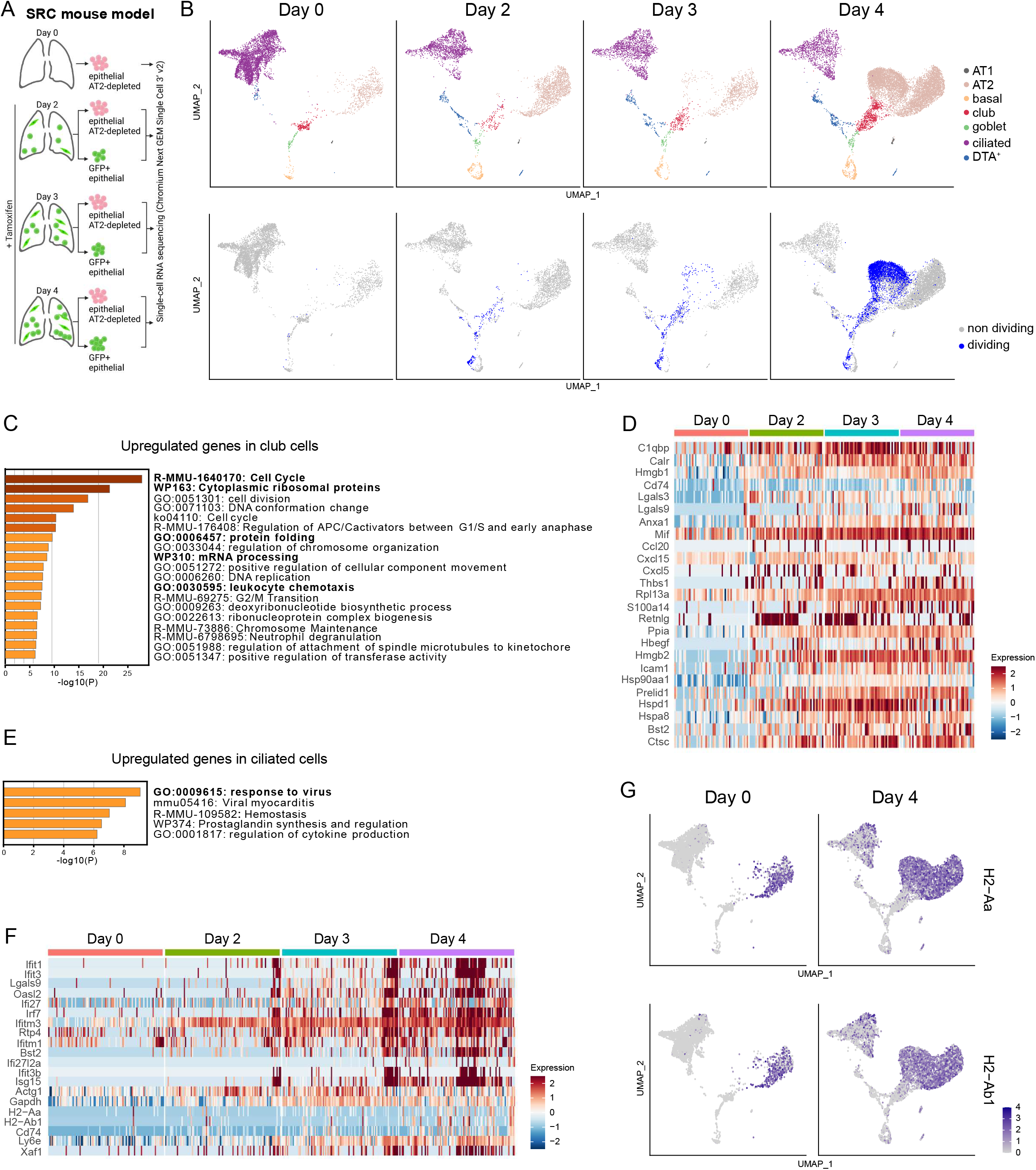
Transcriptional changes in epithelial lung cells after *Scgb1a1\** cell depletion. scRNA-seq analysis was done using lung epithelial cells from SRC mice harvested on day 0 (n=2 mice), day 2 (n=7 mice), day 3 (n=5 mice), and day 4 (n=5). **A.** Outline of cells used for scRNA-seq. Created with BioRender.com. **B.** UMAP embedding showing epithelial cell type and dividing and non-dividing epithelial cells on the indicated days. **C.** Gene set enrichment analysis (GSEA) of upregulated genes in club cells on day 2, 3, and 4 compared to day 0 (FC >2, p_adj <0.05). The terms mentioned in the text are shown in bold. **D.** Heatmap of genes included in the ‘‘leukocyte chemotaxis” cluster of the GSEA in C showing the expression in club cells. **E.** GSEA of upregulated genes in ciliated cells on day 2, 3, and 4 compared to day 0 (FC >2, p_adj <0.05). The terms mentioned in the text are shown in bold. **F.** Heatmap of genes included in the ‘‘response to virus” cluster of the GSEA in E showing the expression in ciliated cells. **G.** UMAP embedding showing the expression of H2-Aa and H2-Ab1 on day 0 and day 4 (p_ adj <0.05 in ciliated cells).

We identified AT1, AT2, basal, club, goblet, ciliated, and DTA^+^ cells (Fig. 3B, Fig. S3B and C, Supplementary Table 3). Basal, club, and goblet cells were again the main cell types dividing early after epithelial injury, while AT2 cells become the dominant dividing cell type as injury progresses (Fig. 3B). We then characterized the transcriptional changes between the cells isolated from uninjured lungs and the cells at the three time points after tamoxifen injection for each cell type (Supplementary Table 4). In goblet, basal, and AT2 cells, the top upregulated DEGs were predominantly associated with cell cycle (Fig. S3E, Supplementary Table 4). While basal cells increased expression of these genes as early as day 2, the expression in AT2 cells did not increase until day 3, which is in accordance with AT2 cells starting to divide later (Supplementary Table 4).

Club cells showed the highest number of DEGs (FC >2, p_adj <0.05) post-injury at all time points, reaching more than 300 DEGs on day 4 (Fig. S3F). Upregulated DEGs were mainly related to cell cycle, cytoplasmic ribosomal proteins, protein folding, mRNA processing, and genes associated with leukocyte chemotaxis (e.g. *Ccl20, Cxcl15, Cxcl5*), the latter suggesting a role of club cells in immune activation (Fig. 3C and D, Supplementary Table 4).

Ciliated cells elevated the expression of interferon-stimulated genes (Fig. 3E and F, Supplementary Table 4), which are involved in viral response mechanisms and are crucial in driving and maintaining lung inflammation, for example, by promoting antigen presentation or stimulating cytokine production^41^. Ciliated cells also upregulated genes implicated in antigen processing and presentation, including MHC class II molecules (MHCII), which are normally found exclusively in professional antigen presenting cells^42^. Unlike AT2 cells, which express high levels of MHCII independent of inflammatory stimuli (Fig. 3G)^43^, ciliated cells demonstrated very low expression of these molecules in homeostasis (Fig. 3G). Thus, the upregulation of MHCII molecules in ciliated cells after epithelial damage suggests that they might function as antigenpresenting cells upon lung epithelial injury.

Together, these data suggest that club and ciliated cells may contribute to the initiation of an inflammatory response following lung injury.

### DTA^+^ cells share distinct gene expression programs and reveal an undescribed lung cell population in COVID-19 patients

Genetic injury mouse models using inducible DTA expression have been widely used since they allow for rapid and controlled cell type-specific ablation^44^. We found that viable DTA^+^ cells can persist in the injured tissue and express a distinct gene expression signature (Fig. 1G), which may provide insights into mechanisms of cellular response to pathological conditions. To characterize these cells in more detail, we further enriched rare cell types and used a more sensitive 3’ RNA sequencing protocol (Fig. 4A, Fig. S4A).

**Fig. 4:**
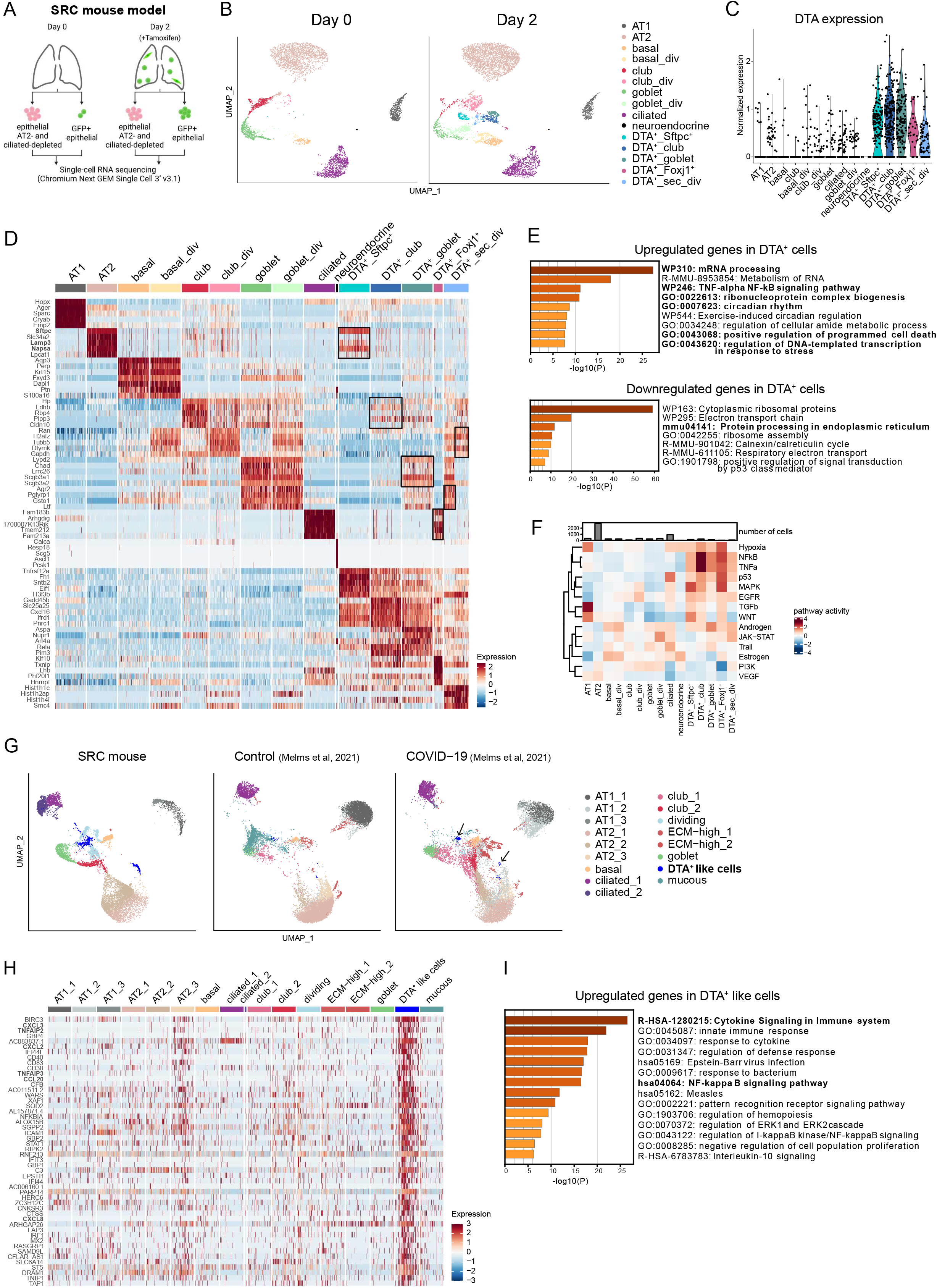
Characterization of DTA^+^ cells and comparison to lung cells from COVID-19 patients. **A.** Outline of cells used for scRNA-seq. Created with BioRender.com. **B.** UMAP embedding showing epithelial cell type assignment before (day 0, n= 8 mice) and after tamoxifen administration (day 2, n=10 mice). **C.** Violin plot with normalized DTA expression in all epithelial cells. **D.** Heatmap of the top five upregulated genes across epithelial cell types on day 2 ranked by power. Scaling of expression was done after downsampling to 100 cells per cell type. Black boxes highlight the expression of the marker genes of the closest epithelial population in DTA^+^ cells. **E.** GSEA of common DEGs in DTA^+^ populations in relation to all non-DTA^+^ cells. The terms mentioned in the text are shown in bold. **F.** Heatmap of the activities of 14 pathways inferred with PROGENY for all epithelial populations on day 2. **G.** UMAP embedding showing cell type assignment of COVID-19 and control human epithelial cells integrated with SRC mouse epithelial cells (day 0 and day 2). Arrows point to DTA-like cells present in the COVID-19 patient samples. **H.** Heatmap of the top 50 DEGs in DTA^+^-like cells from COVID-19 epithelial cells. **I.** GSEA of upregulated genes in the DTA^+^-like cells. The terms in bold depict the clusters that are mentioned in the text.

Unsupervised clustering revealed the same dividing and non-dividing epithelial cell types as described above, neuroendocrine cells, and five clusters of DTA^+^ cells (Fig. 4B and C, Fig. S4B). To determine the cells of origin or the closest relatives of the DTA^+^ populations, we calculated the correlation coefficient of gene expression between the different cell types on day 2 (Fig. S4C). One DTA^+^ cell cluster showed high expression of AT2 canonical marker genes (Fig. 4D). We hypothesized that these cells, which we named DTA^+^_Sftpc^+^, were either derived from a *Scgb1a1^+^* AT2 subpopulation^9^ and could therefore activate DTA expression (Fig. S4D), or from club cells that differentiated into AT2 cells^45^. Two clusters transcriptionally resembled secretory cells and were accordingly named DTA^+^_club and DTA^+^_goblet cells, and DTA^+^ dividing cells included a mixture of cells resembling goblet and club cells, so we refer to them as DTA^+^ secretory dividing (DTA^+^_sec_div) (Fig. 4D and Fig. S4C). The remaining cluster, which we termed DTA^+^_Foxj1^+^, expressed ciliated cell genes (Fig. 4D and Fig. S4C) and likely evolved from club cells that differentiated into ciliated cells^9^. A complete list of the marker genes for each population at day 2 is provided in Supplementary Table 5.

Besides their cell type-specific gene signatures, the DTA^+^ populations shared gene expression patterns and had several marker genes in common (Fig. 4D). The co-upregulated genes showed a prominent enrichment in mRNA processing genes, including mRNA splicing-related genes (Fig. 4E, Supplementary Table 6). In addition, DTA^+^ populations displayed upregulation of genes involved in ribonucleoprotein complex biogenesis, cellular response to stress, induction of apoptosis, and downregulation of protein processing-related genes (Fig. 4E, Supplementary Table 6), which could be a consequence of DTA-mediated protein synthesis inhibition^46^. Curiously, all DTA^+^ populations, apart from DTA^+^_Foxj1^+^ cells, displayed increased levels of *Cftr*, a transmembrane conductance regulator that is mutated in cystic fibrosis (Fig. S4E). This gene is highly expressed by ionocytes found in the proximal airways of human and mice^11,29^. However, DTA^+^ cells did not express other ionocyte marker genes, implying that these are different cell types (Fig. S4E). Finally, DTA^+^ cells showed an enrichment in genes involved in pro-inflammatory TNF-alpha/NF-κB signaling and circadian rhythm, which could influence the inflammatory response^47^ (Fig. 4E, Supplementary Table 6). A complementary pathway activity analysis based on core pathway responsive genes confirmed that DTA^+^ populations had increased activation of several inflammatory pathways such as TNF, NF-κB, MAPK, and EGFR (Fig. 4F).

Considering these specific transcriptional changes in DTA^+^ cells, we hypothesized that they might mirror characteristics of damaged cells in the context of human lung diseases. Using gene signatures from DisGeNET^48^ and MSigDB^49, 50^ databases for several human lung diseases, we found that DTA^+^ cells had particularly high enrichment for genes expressed in epithelial lung cells from COVID-19 patients^51^ (Fig. S4F). Integration of our murine dataset with a scRNA-seq dataset from a COVID-19 study^4^ revealed a new epithelial cell population unique to COVID-19 samples (Fig. 4G). This cell population showed differential upregulation of genes involved in TNF and NF-κB signaling, supporting the similarity to DTA^+^ cells (Fig. 4H and I, Supplementary Table 7).

Together, epithelial cells expressing DTA can persist in the lung and share a distinct gene expression profile that exhibits activation of inflammatory signaling pathways and remarkable transcriptional similarity to a previously undescribed lung cell population in patients with COVID-19.

### DTA^+^ cells signal to immune, epithelial, and mesenchymal cells

Inflammatory pathways, such as the ones described above, can be activated in epithelial cells by microbial components and lead to the initiation of an immune response through chemokine secretion^52^. Upregulated genes in the COVID-19 DTA^+^-like population were enriched for genes related to cytokine signaling in the immune system (Fig. 4I), and several chemokines were among the top 50 DEGs (Fig. 4H), suggesting that the identified COVID-19 cell population can contribute to the inflammatory response in these patients. To investigate this further, we used the murine DTA^+^ cells as model for the human COVID-19 cell population and investigated their potential role in immune cell activation with focus on chemokines.

After *Scgb1a1^+^* cell depletion, chemokine expression increased only slightly in regular epithelial cells, with the exception of *Ccl20* in club cells and *Cxcl17* in club_div and ciliated cells (Fig. 5A and S5A). In contrast, DTA^+^ cells were a major source of chemokine production, and DTA^+^_club and DTA^+^_Foxj1^+^ even dedicated a large proportion of their total transcriptome to chemokine production (2% in DTA^+^_club at day 2 vs. 0.7% in club cells at day 0; 0.4% in DTA^+^_Foxj1^+^ at day 2 vs. 0.1% in ciliated cells at day 0) (Fig. 5A, Fig. S5B). Among the chemokines significantly upregulated in at least one DTA^+^ cell type, but not in any regular epithelial cell type, were *Ccl9, Ccl20, Cxcl1, Cxcl2, Cxcl3, Cxcl10*, and *Cxcl16* (Fig. 5B). Ccl9, *Cxcl2*, and *Cxcl16* were shown to be expressed in immune cells during homeostasis, whereas *Ccl20, Cxcl1, Cxcl3*, and *Cxcl10* are barely expressed in any cell type of the lung under healthy conditions (Fig. S5C)^53^, suggesting that they are specifically produced under damage-inducing conditions. These DTA^+^-specific chemokines are known to recruit neutrophils (CXCL1, CXCL3), dendritic cells (CCL20), T cells (CCL20, CXCL10), and B cells (CCL20)^54^.

**Fig. 5:**
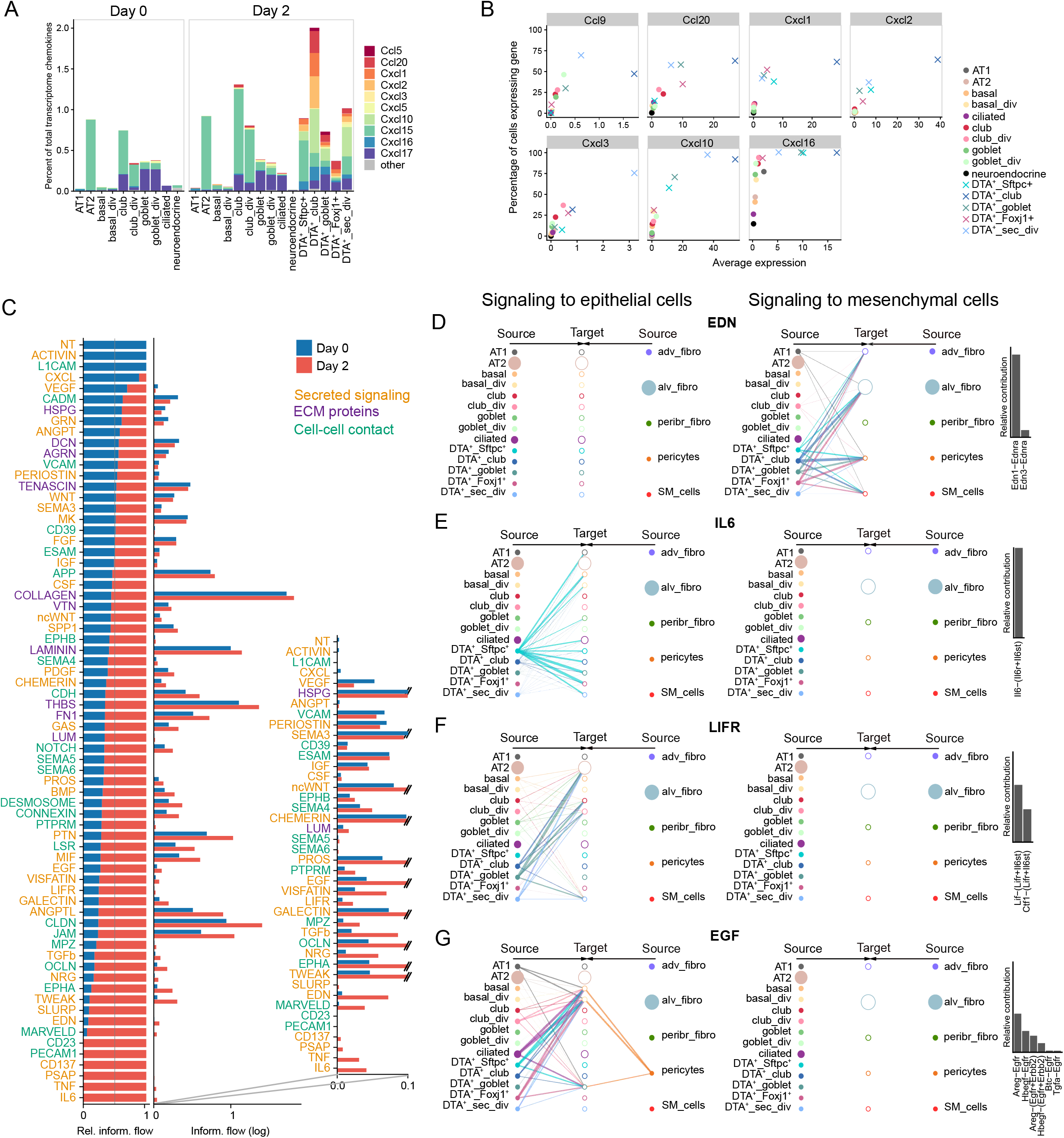
Chemokine expression and crosstalk to mesenchymal and epithelial cells of DTA^+^ cells. **A.** Chemokine expression in epithelial lung cells from SRC mice before (day 0) and after injury (day 2). Genes with a mean expression of at least three counts per 10,000 in any cell type are colored. **B.** Expression of chemokines upregulated in DTA^+^ epithelial cells after injury (FC >2 and p_adj <0.05 in any DTA^+^ cell type). The percentage of cells expressing the gene are plotted against the average expression per cell type. **C.** Ranking of significant signaling pathways based on differences of overall information flow (i.e. summed interaction strengths) within the inferred networks on day 0 and day 2. Left: Relative information flow. Middle: Absolute information flow (log scale). Right: Zoom in on the absolute information flow of pathways with log(information flow) < 0.1 at any time point. Signaling pathways are colored by the type of interaction (yellow: secreted signaling, purple: ECM-receptor, green: cell-cell contact). **D-G.** Hierarchical plots showing the inferred communication network for EDN **(D)**, IL6 **(E)**, LIFR **(F),** and EGF **(G)** signaling. Circle sizes are proportional to the number of cells in each cell type, and line width corresponds to the interaction strength. The relative contributions of ligand-receptor pairs making up at least 1% of the pathway strength are shown on the right.

To characterize the interactions between epithelial and mesenchymal cells, we analyzed the ligand-receptor expression of all epithelial and mesenchymal cells in lungs of SRC mice at day 0 and day 2. After injury, new or increased signal transmission occurred in several pathways (Fig. 5C), mainly as a result of new ligand-receptor pairings involving DTA^+^ epithelial cells (Fig. S5D and E). Specifically, signaling networks involved in the formation of tight junctions such as MARVELD, OCLN, JAM, and CLDN were activated in epithelial cells, with particularly high levels in DTA^+^ cell types (Fig. 5C, Fig. S5F). Tight junctions are crucial during lung repair as they are responsible for sealing the epithelial barrier and preventing the entry of pathogens and the free diffusion of solutes into airspace^55^. *Edn1* was also specifically expressed in DTA^+^ epithelial cells and was predicted to signal to alveolar fibroblasts, adventitial fibroblasts, smooth muscle cells, and pericytes (Fig. 5D, S5F and G). EDN1 was shown to stimulate fibroblast replication, migration, and collagen synthesis^56^, suggesting a role of DTA^+^ cells in the activation of mesenchymal cells after epithelial injury. Finally, we found that DTA^+^ cells highly expressed several ligands that can signal to other epithelial cells and support lung regeneration (Fig. 5E-G, Fig. S5F and G). Specifically, Il6, predicted to signal from DTA^+^_Sftpc^+^ cells to AT1, basal, goblet, and ciliated cells, was found to be crucial for lung repair after influenza-induced lung injury in mice^57^. LIFR pathway, which was previously shown to protect lung tissue during pneumonia in mice^58^, was predicted to be activated in AT2 and club cells by DTA^+^ cells through the expression of LIF. DTA^+^ cells also expressed high levels of *Areg* and *Hbegf* that can signal to basal cells through EGFR, previously shown to be essential for basal cell proliferation^59^.

Overall, DTA^+^ cells express specific chemokines and growth factors that can trigger an immune response and contribute to lung regeneration through epithelial and mesenchymal cell support.

### Combined depletion of *Scgb1a1^+^* and *Sftpc^+^* cells allowed the identification of basal *Krt13^+^* and club *Krt15^+^* in the mouse distal lung

We next characterized the consequences of increased epithelial damage in the peripheral lung through additional depletion of AT2 cells. Therefor, we crossed SRC mice with mice expressing tamoxifen-inducible Cre recombinase in AT2 cells under the *Sftpc* promoter^60^. The resulting Sftpc-CreER x Scgb1a1-CreER x Rosa26R-DTA x CycB1-GFP (SfSRC) mice (Fig. 6A) allow simultaneous depletion of *Scgb1a1^+^* and *Sftpc^+^* cells after tamoxifen administration, and tracing of GFP^+^ dividing cells (Fig. 6A). *Scgb1a1^+^* and *Sftpc^+^* cells progressively disappeared from day 2 to 14, and numerous dividing cells emerged in the airways, underlying tissue, and alveoli (Fig. 6B). Airway regeneration was not complete at day 28 based on cell morphology and Cyp2f2 club cell staining, despite only few dividing cells remaining, but was finally achieved at day 56. Lung morphology on days 14, 28, and 56 showed no evidence of fibrosis (Fig. 6B), and Masson staining at day 14 confirmed the absence of collagen deposition (Fig. S6A).

**Fig. 6:**
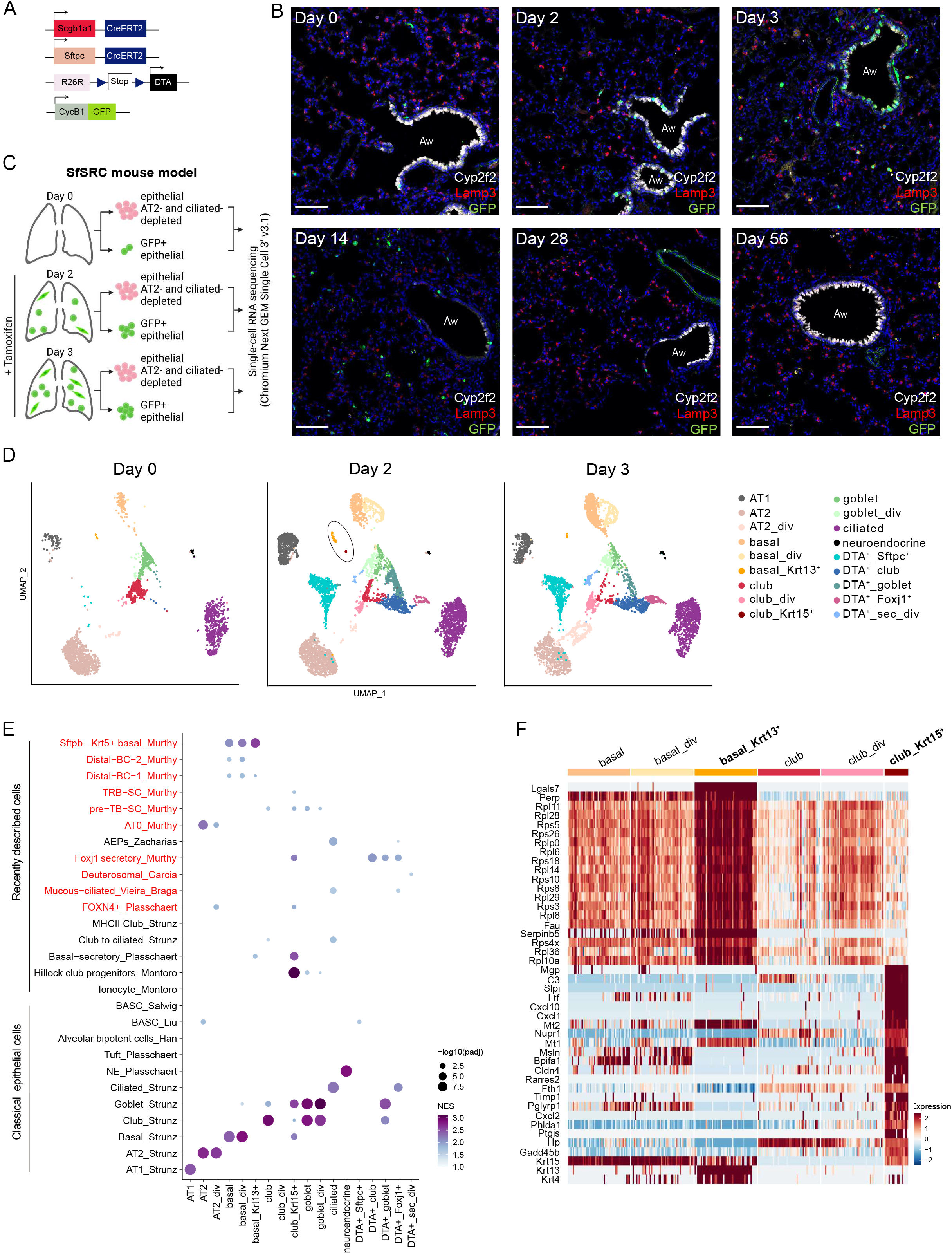
Characterization of lung epithelial cells in the SfSRC mouse model. **A.** Schematic of Sftpc-CreER, Scgb1a1-CreER, Rosa26R-DTA, and CycB1-GFP transgenes in SfSRC mice. **B.** IF staining of SfSRC mouse lungs with Cyp2f2 (white; club cells), Lamp3 (red; AT2 cells), CycB1-GFP (green; dividing cells), and DAPI (blue; nuclei), on days 0, 2, 3, 14, 28, and 56 after tamoxifen injections. Aw: airway. Scale bar: 100 μm. Images are representative of at least three animals analyzed per group. **C.** Outline of cells used for scRNA-seq. Created with BioRender.com. **D.** UMAP embedding showing cell type assignment of epithelial cells before (day 0, n=5 mice) and two (n=9 mice) and three days (n=6 mice) after tamoxifen injection. At day 2, basal_Krt13^+^ and club_Krt15^+^ cells are highlighted. **E.** Dot plot showing normalized enrichment score (NES) and significance for published signatures of epithelial cell types in all SfSRC mouse epithelial populations (all timepoints merged). DATP: damage-associated transient progenitors; PATS: pre-alveolar type-1 transitional cell state; RAS: respiratory airway secretory; BC: basal cell; TRB-SC: terminal and respiratory bronchioles secretory cell; pre-TB-SC: pre-terminal bronchiole secretory cell; AEP: alveolar epithelial progenitor; BASC: bronchioalveolar stem cell. Cell types described in the human lung are depicted in red. **F.** Heatmap of the top 20 marker genes for basal_Krt13^+^ and club_Krt15^+^ cells, and expression of Krt15, Krt13, and Krt4, in basal and club cells of SfSRC mice (all timepoints merged).

We analyzed lung epithelial cells from SfSCR mice with scRNA-seq before (day 0) and 2 and 3 days after tamoxifen injection (Fig. 6C). Cells were sorted analogously to the SRC model (Fig. S4A) to enrich rare EPCAM^+^ and dividing GFP^+^ cells (Fig. S6B). We were able to distinguish all previously identified cell types and two additional small clusters (Fig. 6D, Fig. S6C-E, Supplementary Table 8). One cluster comprised *Krt13* basal cells and the other *Krt15^+^* club cells (Fig. 6D). Projection of cells from SfSRC mice onto cells from SRC mice and integration of all datasets generated in this study showed that *Krt13* basal cells were present in both models, while *Krt15^+^* club cells were unique to the SfSRC model (Fig. S6F and G).

To systematically assess the similarity between the two new cell populations and reported cell types, we performed fast gene set enrichment analysis (FGSEA) with gene signatures of canonical lung epithelial cell types and recently described new lung epithelial cells^7,11,13,14,29,31, 53, 61,62, 63^. Although *Krt13^+^* basal cells were previously described in the mouse trachea^11^, our *Krt13^+^* basal cells were transcriptionally distinct (Fig. 6E). Instead, they closely resembled *Sftpc^-^ Krt5^+^* basal cells found in human airways (Fig. 6E and F). The *Krt15^+^* club cell cluster was transcriptionally similar to hillock club cells, despite the fact that they did not consistently express the main marker genes *Krt4* and *Krt13* (Fig. 6E and F). Since hillock club cells were found exclusively in the trachea^29^, and we removed trachea and main bronchi before performing scRNA-seq, we hypothesized that hillock-like cells can be present in the distal airways.

Overall, the combined depletion of *Scgb1a1^+^* and *Sftpc^+^* cells revealed two new cell types, basal *Krt13^+^* that resembled human *Sftpc^-^ Krt5^+^* basal cells, and a population of hillock-like club cells expressing *Krt15^+^*.

### Trajectory modeling predicts differentiation of goblet cells into basal and club cells

Finally, we explored the relations and predicted biological trajectories between the identified epithelial cell types in response to *Scgb1a1^+^* and *Sftpc^+^* cell loss. To get a better understanding of the data topology in the SfSRC model, we visualized the cells before and after injury in a force-directed graph (Fig. 7A) and summarized the connectivities with partition-based graph abstraction (PAGA) (Fig. 7B). The edges in the PAGA graph represent possible differentiation paths, which we analyzed using diffusion maps of different cell type subsets to obtain finer resolution.

**Fig. 7:**
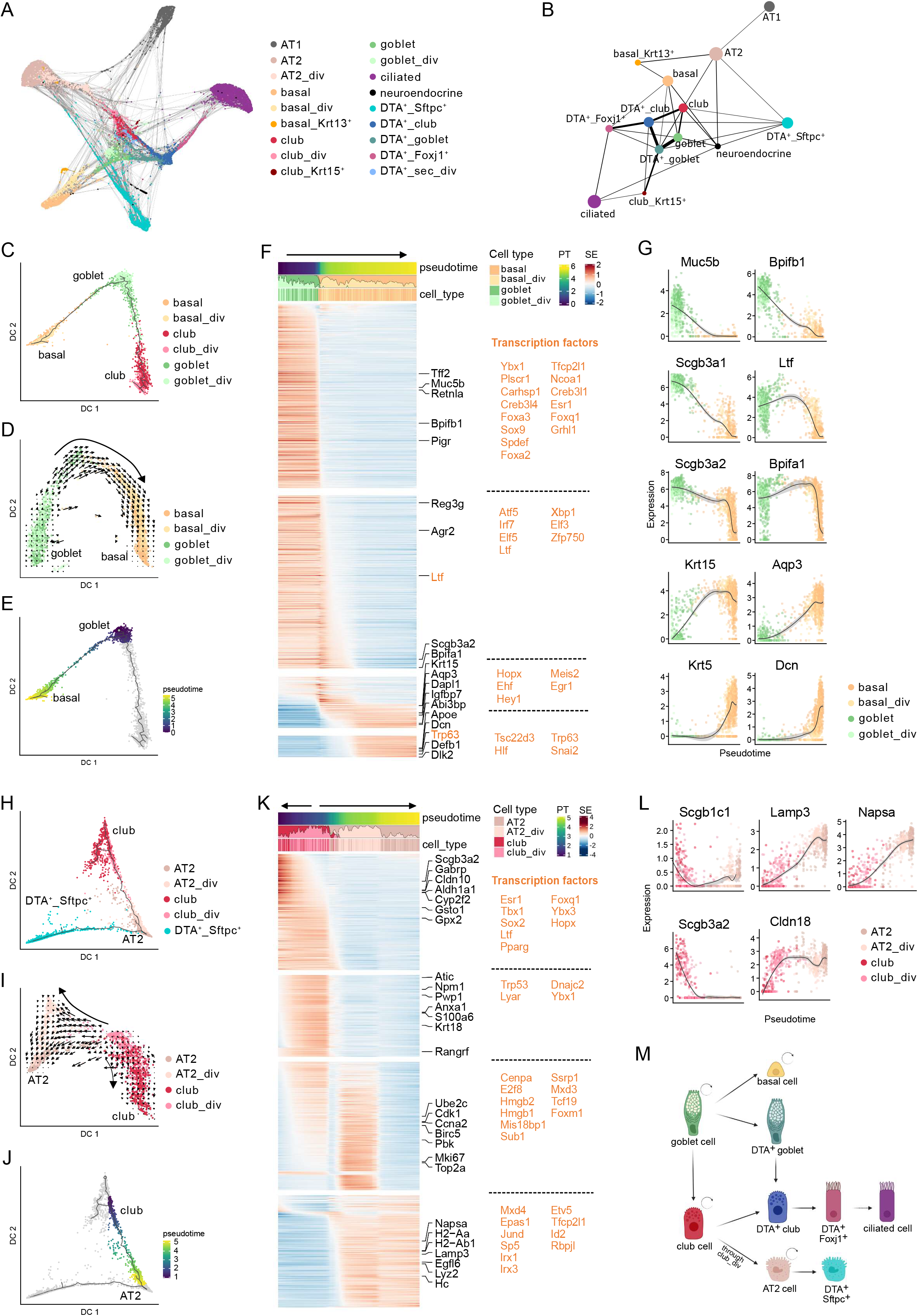
Differentiation trajectories between epithelial lung cells in SfSRC mice. **A.** Force-directed graph of single cells with edges to the 10 nearest neighbors. **B.** PAGA graph summarizing the connectivities between different cell types. The thickness of the lines is a statistical measure of connectivity between cell types. Dividing cell types were excluded for simplicity. **C.** Diffusion map of dividing and non-dividing basal, goblet, and club cells with pseudotime trajectory colored by cell type. **D.** Diffusion map of dividing and non-dividing basal and goblet cells with RNA velocity vectors indicating differentiation of goblet into basal cells. **E.** Diffusion map of dividing and non-dividing basal and goblet cells with pseudotime of cells selected to characterize the goblet to basal cell trajectory in F. **F.** Smoothed expression heatmap of the top 1,000 altered genes along the differentiation trajectory from goblet to basal cells. The names of the top five genes in each cluster are annotated and transcription factors are indicated in orange. All transcription factors of each cluster are listed on the right. PT: pseudotime; SE: scaled expression. **G.** Normalized expression along the goblet to basal cell pseudotime trajectory for exemplary genes. The black line is the smoothed expression with the confidence interval shown in gray. **H.** Diffusion map of dividing and non-dividing club, AT2, and DTA^+^_Sftpc^+^ cells with pseudotime trajectory colored by cell type. **I**. Diffusion map of dividing and non-dividing club and AT2 cells with RNA velocity vectors indicating that dividing club cells give rise to club cells and AT2 cells. **J.** Diffusion map of dividing and non-dividing club and AT2 cells and DTA^+^_Sftpc^+^ cells with pseudotime trajectory colored by pseudotime of the cells selected to characterize the club to AT2 cell trajectory in K. **K.** Smoothed expression heatmap of the top 1,000 altered genes along the differentiation trajectory from club to AT2 cells (see Fig. 7J). **L.** Normalized expression along the club to AT2 cell pseudotime trajectory for exemplary genes. Figures were calculated on merged data from all timepoints (day 0, 2, and 3). **M.** Proposed model of differentiation routes (straight arrows) and self-renewal capacities (curved arrows) of lung epithelial cells based on data and mouse models in this study. Created with BioRender.com.

According to the basic lineage model of the lung epithelium, basal cells are considered to be progenitors of club cells, which in turn give rise to goblet and ciliated cells^32^. However, in the PAGA map, basal cells were strongly connected to goblet cells and not to club cells (Fig. 7B). A diffusion map restricted to basal, goblet, and club cells confirmed that goblet cells serve as bridge between basal and club cells (Fig. 7C). The directionality inferred by RNA velocity even suggests that goblet cells give rise to basal cells (Fig. 7D and S7A), which is in accordance with a recently described dedifferentiation trajectory from goblet to basal cells following bleomycin lung injury^64^. We modeled a pseudotime trajectory from the tip of the goblet cell cluster and calculated DEGs along the goblet-basal and goblet-club axes separately. The transition from goblet to basal cells is primed by a decrease of the goblet cell markers *Muc5b, Bpifb1, Scgb3a1, Scgb3a2*, and *Bpifa1*, and an increase of the basal cell markers *Aqp3, Krt15, Krt5*, and *Dcn* (Fig. 7E-G, Supplementary Table 9).

Next, we followed the goblet cell trajectory to club cells (Fig. S7B-E). The key transcription factors required for goblet cell differentiation and mucus production *Spdef* and *Foxa3*^65^ are among the first downregulated genes, followed by goblet cell markers including *Agr2, Bpfib1, Muc5b*, and *Scgb3a1* (Fig. S7D-E, Supplementary Table 10). At the same time, the expression of club cell markers such as *Cckar, Scgb1c1*, and *Sftpb* increased. As in the goblet-basal cell trajectory, the differentiation between goblet and club cell clusters occurs without proliferation (Fig.7D and Fig. S7B). PAGA and diffusion maps also showed a strong connection between DTA^+^_goblet and DTA^+^_club cells, suggesting that DTA^+^_goblet cells differentiate into club cells similar to uninjured goblet cells (Fig. 7B and Fig. S7F).

Club cells are known to give rise to ciliated cells^9^. We also observed a trajectory from club to ciliated cells, which occurred almost exclusively through DTA^+^_club (Fig. S7G and H), suggesting that differentiation into ciliated cells occurs preferentially through stressed club cells. While the expression of club cell markers is gradually reduced during the transition to DTA^+^_club and completely lost in DTA^+^_Foxj1^+^ cells, the expression of ciliated markers already started to increase in DTA^+^_club (Fig. S7I-K).

Finally, club_div cells were predicted to give rise to both club and AT2 cells (Fig. 7H and I). As observed in SRC mice, club_div cells had decreased secretory cell markers, such as *Scgb3a2* and *Scgb1c1*, and increased AT2 markers, such as *Napsa* and *Lamp3* (Fig. 7J-L, Supplementary Table 12). *Cldn18*, a tight junction protein highly expressed in the alveolar epithelium, was also among the top DEGs upregulated in club_div, corroborating the club to AT2 cell transition (Fig. 7L).

Taken together, these data suggest secretory cells as the main epithelial progenitor cell population in the distal lung following *Scgb1a1^+^* and *Sftpc^+^* cell loss. In particular, goblet cells are predicted to be progenitors for basal and club cells, and club_div cells for club and AT2 cells. Stressed (DTA^+^) club cells differentiate into ciliated cells (Fig. 7M).

## DISCUSSION

This study provides a detailed single-cell transcriptome analysis of the lung epithelial and mesenchymal responses to targeted epithelial damage using novel genetically-modified mouse models and cell sorting strategies to enrich for rare and dividing cells. We identify and characterize previously undescribed cell types and provide new insights into functional roles of lung epithelial cells and their differentiation paths.

Elimination of *Scgb1a1^+^* cells or both *Scgb1a1^+^* and *Sftpc^+^* cells resulted in immediate proliferation of most epithelial cell types with the exception of ciliated and AT1 cells known to be terminally differentiated. In particular, AT2 cells showed a remarkable increase in cell division even when only *Scgb1a1^+^* club cells were targeted, which is unexpected since AT2 cells do not contribute to bronchiolar epithelium regeneration. The reason for the pronounced AT2 cell proliferation is unclear but could be related to an indirect role of AT2 cells in lung regeneration. For example, a subset of AT2 cells was recently shown to play a critical role in maintaining barrier immunity in the lung by regulating the function of memory CD4^+^ T cells^66^.

Based on our idea that the use of chemical or infectious agents could destroy yet unknown cells involved in regeneration, one of our goals was to search for new cell types through the application of targeted genetic cell disruption. Indeed, our study revealed rare cell populations that we named basal-goblet_div, club_div, *basal_Krt13^+^*, and *club_Krt15^+^* cells, which were not previously described in the mouse distal lung. Club_div cells did not display gene expressing signatures of known club progenitor cells but were predicted to be the source of club and AT2 cells upon *Scgb1a1^+^* and *Sftpc^+^* cell depletion. Likewise, basal-goblet_div cells, expressing both basal and goblet cell marker genes, were never described in the mouse or human respiratory tract and, given their proliferative response, could represent a new progenitor population. Conversely, *basal_Krt13^+^* cells are closer to human *Sftpc^-^ Krt5^+^* basal cells than to mouse basal cells. Finally, *club_Krt15^+^* cells displayed a transcriptional profile similar to tracheal hillock cells, suggesting their presence in the distal mouse lung. Further analyses are required to validate the presence, precise location, and function of these newly identified cells.

Goblet cells are the main mucus-producing cells in the airways and are thought to be terminally differentiated cells with origin in club cells^32^. We found that goblet cells are capable of dividing and trajectory modelling indicates that they differentiate into club and basal cells instead of the other way around. The latter observation is supported by a single-cell transcriptomics study that predicted dedifferentiation from goblet to basal cells in a bleomycin injury mouse model^64^. Furthermore, Tata *et al*. showed that basal cell ablation using Krt5^+^-DTA mice induced dedifferentiation of secretory cells into basal cells by *in vivo* lineage tracing of *Scgb1a1^+^* cells^17^, a marker shared by club and goblet cells^63^. The authors also demonstrated that cells expressing *Atp6v1b1*, a marker for goblet cells in our study (Supplementary Table 5 and Supplementary Table 8), dedifferentiated into basal cells in an organoid culture system. Together, these findings strongly suggest that goblet cells can function as progenitor cells in the distal mouse lung and actively contribute to epithelial repair.

Severe COVID-19 is characterized by persistent lung inflammation that can lead to the development of acute respiratory distress syndrome and death, but it is not clear what triggers inflammation^67^. Recently, it has been shown that, in COVID-19 patients, SARS-CoV-2 can infect peripheral blood monocytes and tissue-resident macrophages, and activate an inflammatory cascade with a unique transcriptome that results in inflammatory cell death^68, 69^ It was suggested by the authors that the release of inflammatory factors upon cell death can be a trigger for the overactive immune inflammatory response seen in COVID-19. In our study, the transcriptional profile of persistent DTA^+^ cells in the lungs allowed us to identify a subpopulation of lung epithelial cells from COVID-19 patients, as they share increased expression of key inflammatory factors and pathways. The presence of this population in the lungs of COVID-19 patients suggests that epithelial cells can also be a source of immune activation in these patients. Further characterization of the immune response in injured lungs of SRC and SfSRC mice and the newly identified cell population in SARS-CoV-2-infected patient lungs may help to better understand the overshooting immune response in COVID-19 patients and identify crucial interactions between immune and epithelial cells during lung regeneration.

Overall, we provide valuable new insights into the cellular and molecular response of lung cells to pathological conditions and epithelial regeneration mechanisms. In addition, we contribute with models and a comprehensive resource that can form the basis for further studies, particularly functional studies.

## METHODS

### Mouse experiments

Mice were housed within the pathogen-free DKFZ animal facility. All experiments were approved by the regional authority in Karlsruhe, Germany, under protocol numbers G-238/14 and G-303/19 and performed according to federal and institutional guidelines. Scgb1a1-CreER, Rosa26R-DTA, and CycB1-GFP mice were purchased from The Jackson Laboratory and used to generate the SRC mouse line. SPCcreT2rtTA mice^60^ (Sftpc-CreER), kindly provided by Rocio Sotillo (DKFZ, Heidelberg, Germany), were crossed with SRC mice to generate the SfSRC mouse line. The mice were homozygous for all transgenes except CycB1-GFP, which had multiple transgenes integrated in the genome.

Mice were injected intraperitoneally with 4 mg tamoxifen dissolved in corn oil once (Fig. 1, S1, 2 and S2) or on two consecutive days (all other figures). Control animals (day 0) were not injected. Analysis was done at the indicated timepoints, counting from the first day of tamoxifen injection. Mice were sacrificed by cervical dislocation, the abdominal aorta and vena cava were severed, and lungs were perfused with PBS though the right ventricle. Further processing of the lungs depended on the type of downstream analysis.

### Immunofluorescence

Lungs were carefully inflated with 1% PFA (Thermo Fischer Scientific) though the trachea, fixed overnight at 4°C, washed in PBS, subjected to a 10-30% sucrose gradient, embedded in OCT compound (Sakura), and snap-frozen in ethanol/dry ice. Cryosections with 10 μm thickness were placed on SuperFrost UltraPlus Gold adhesion slides (Menzel) and stained using immunostaining chambers (Thermo Fischer Scientific). Briefly, sections were washed with PBS and incubated with a perm/block solution containing 0.3% Triton X-100 (Sigma) and 5% BSA (Sigma) in PBS for 30 minutes at room temperature (RT). Primary antibodies were incubated at RT for 60 minutes, sections were washed with 0.1% Tween 20 (Sigma) in PBS, and secondary antibodies were incubated for 30 minutes at RT followed by washing with 0.1% Tween 20 in PBS. The following antibodies were used: acetylated tubulin (1:1000, #T7451, Sigma), Agr2 (1:100, #12275-1-AP, Proteintech), CC10 (1:500, #07-623, Millipore), CD34 (1:200, #553731, BD), Cyp2f2-Alexa Fluor 647 (1:1000, #sc-374540, Santa Cruz), GFP-FITC (1:500, #600-102-215, Rockland), Krt5 (1:500, # ab53121, Abcam), Lamp3 (1:500, #DDX0192A647-100, Origene), Npnt (1:50, PA547610, Thermo Fischer Scientific), Pdpn (1:2000, #ab11936, Abcam), SPC (1:1000, # AB3786, Millipore), donkey anti-rabbit Alexa Fluor 568 (1:500, Thermo Fischer Scientific), goat anti-rat Alexa Fluor 568 (1:500, Thermo Fischer Scientific), and goat anti-hamster Alexa Fluor 647 (1:500, Thermo Fischer Scientific). The acetylated tubulin antibody was conjugated with Pacific Blue Antibody Labelling Kit (Thermo Fisher Scientific) prior to use. All antibodies were diluted using the perm/block solution. When indicated, sections were incubated with 0.6μg/ml of DAPI solution (BD Biosciences) and washed with PBS before mounting with Prolong Diamond Antifade mounting media (Thermo Fischer Scientific). Images were acquired using the A1R confocal microscope (Nikon) and processed using Fiji^70^.

### Masson staining

PBS-perfused lungs were inflated with 1% PFA (Thermo Fischer Scientific) and fixed for 24h at 4°C. Lungs were dehydrated in increasing concentration of ethanol and embedded in paraffin. Sections of 3 μm were placed on SuperFrost UltraPlus Gold adhesion slides (Menzel), deparaffinized with histoclear (Linaris), and hydrated in decreasing concentrations of ethanol before staining. Staining was performed with kit trichrome de Masson (RAL diagnostics), Anilin Blue variation, according to the manufacturer’s instructions. Briefly, sections were incubated in Mayer Haemalum for 10 minutes, rinsed in running tap water, stained with Ponceau Fuchsin for 5 minutes, rinsed in 2 baths of 1% acetic water, dipped in phosphomolybdique, stained in Aniline blue for 5 minutes, and rinsed in 2 baths of 1% acetic water. Sections were dehydrated and dipped in histoclear before mounting with Entellan (Millipore). Images were acquired using the Ni-E microscope (Nikon) and processed using Fiji^70^.

### Generation of lung single-cell suspensions

PBS-perfused lungs were inflated with 2 ml of digestion cocktail containing 50 U/ml dispase (Corning), 250 U/ml collagenase type I (Worthington), 5 U/ml elastase (Worthington), and 60 U/ml DNAse I (Roche). Trachea was clipped distally and lungs were dissected in a petri dish on ice to remove extrapulmonary airways (trachea and main bronchi). Lung lobes were placed in a C tube (Miltenyi) containing 3 ml of digestion cocktail, and the m_lung_01 program was run on gentleMACS (Miltenyi). C tubes were placed in a rotating incubation oven at 37°C for 30 minutes. The m_lung_02 program was run again, and the tubes were placed on ice for the next steps. The lung cells were passed through a 70 μm cell strainer (Corning) and centrifuged at 400 g for 5 minutes. The pellet was resuspended in a red blood lysis buffer solution (0.15 M NH4Cl, 10 mM KHCO3, 0.1 mM EDTA), incubated for 2 minutes on ice, and washed with EasySep buffer (STEMCELL Technologies) at 400 g for 5 minutes. To isolate mesenchymal cells (Fig. 2 and S2), a milder digestion cocktail was used containing only 375 U/ml collagenase and 60 U/ml DNase I.

### Fluorescence-activated cell sorting

Lung single-cell suspensions were resuspended in EasySep buffer, incubated with primary antibodies for 20 minutes at 4°C, washed with EasySep buffer at 400 g for 5 minutes, and, in the case of unconjugated primary antibodies, secondary antibodies were added, incubated for 15 minutes at 4°C and washed again. The following antibodies were used, according to the cell populations sorted: CD31 APC (1:200, BD Biosciences), CD45 APC (1:200, BD Biosciences), Ter119 APC (1:200, Life Technologies), Epcam PECy7 (1:300, BD Biosciences), CD24 PE (1:200, BD Biosciences), MHCII Alexa Fluor 700 (1:200, Invitrogen), Npnt (1:40, Invitrogen), Pdgfra BV711 (1:100, BD Biosciences), Sca-1 APC-Cy7 (1:200, BD Biosciences), CD34 PerCP5.5 (1:100, BD Biosciences), and donkey anti-goat Alexa Fluor 680 (1:500, Invitrogen). DAPI at 0.6 ug/ml (BD Biosciences) was added before acquisition to exclude dead cells. Hematopoietic (CD45^+^ and Ter119^+^) and endothelial (CD31^+^) cells were excluded. Dividing cells were sorted based on their expression of GFP. For mesenchymal cell analysis, GFP^+^ dividing cells were run on Chromium as separate samples. For epithelial cell analysis, dividing GFP^+^ Epcam^+^ cells were sorted and added to the Epcam^+^ cells before running the samples on Chromium. To enable enrichment of infrequent cell types, AT2 cells were partially depleted using an anti-MHC II antibody^71^ or AT2 and ciliated cells were partially depleted using a CD24 antibody. Gating strategies are depicted for each experiment. Sorting was performed with a BD FACS Aria II (BD Biosciences) with a 100 μm nozzle.

### Lung organoid cultures

Lung single-cell suspensions from non-injured SRC lungs were sorted according to the sorting strategy described in Figure S2. Approximately 1,000 progenitor epithelial cells (DAPI^-^ CD45^-^CD31^-^ Ter119^-^ Epcam^high^ CD24^dim^)^34^ were resuspended with approximately 20,000 adventitial fibroblasts (DAPI^-^ CD45^-^ CD31^-^ Ter119^-^ Epcam^-^ CD34^+^ Sca-1^+^) or the same number of alveolar fibroblasts (DAPI^-^ CD45^-^ CD31^-^ Ter119^-^ Epcam^-^ CD34^-^ Sca-1^-^ Pdgfra^+^ Npnt^+^) in 50% Growth Factor Reduced Matrigel (Corning). Cell suspensions were carefully dispensed into 0.4 μm pore polyethylene terephthalate Falcon cell culture inserts (Corning), placed in 24-well plates with DMEM-F12 medium supplemented with 5% FBS, 1% insulin/transferrin/selenium (Thermo Fisher Scientific), and 0.05 μg/ml FGF10 (GenScript). 0.1% ROCK inhibitor (Sigma-Aldrich) was added for the first 48h of culture. Medium was changed every 2-3 days and organoids were cultured for 3-4 weeks. Imaging of the organoids was performed with the Lionheart FX Automated Microscope (BioTek) by performing tiles and z-stack images, and the Gen5 3.05 software was used for image stitching and z-stack projections.

### Mass spectrometry

#### Sample preparation

90,000-140,000 adventitial fibroblasts (DAPI^-^ CD45^-^ CD31^-^ Ter119^-^ Epcam^-^ Pdgfra^+^ CD34^+^ Sca-1^+^), 200,000 alveolar fibroblasts (DAPI^-^ CD45^-^ CD31^-^ Ter119^-^ Epcam^-^ CD34^-^ Sca-1^-^ Pdgfra^+^), and 160,000-230,000 Pdgfra^-^ cells (DAPI^-^ CD45^-^ CD31^-^ Ter119^-^ Epcam^-^ CD34^-^ Sca-1^-^ Pdgfra^-^) were sorted from uninjured lungs in two independent experiments (n=6 and n=4 mice, respectively), and lysed in RIPA buffer (Thermo Fisher Scientific) supplemented with protease inhibitor cocktail (Roche). Further sample preparation and data analysis was performed by the DKFZ Genomics and Proteomics Core Facility as follows: SDS-PAGE gel-based protein purification was performed before trypsin digestion of the proteins on a DigestPro MSi robotic system (INTAVIS Bioanalytical Instruments AG) according to an adapted protocol by Shevchenko *et al*.^72^. Peptides were separated on a cartridge trap column, packed with Acclaim PepMap300 C18, 5 μm, 300 Å wide pore (Thermo Fisher Scientific) in a three step, 180 min gradient from 3% to 40% ACN on a nanoEase MZ Peptide analytical column (300 Å, 1.7 μm, 75 μm x 200 mm, Waters) carried out on a UltiMate 3000 UHPLC system. Eluting peptides were analyzed online by a coupled Q-Exactive-HF-X mass spectrometer (Thermo Fisher Scientific) running in data depend acquisition mode, where one full scan at 120 k resolution (375-1500 m/z,3e6 AGC tagert, 54 ms maxIT) was followed by up to 35 MSMS scans at 15 k resolution (1e5 AGC tagert, 22 ms maxIT) of eluting peptides at an isolation window of 1.6 m/z and a collision energy of 27% NCE. Unassigned and singly charged peptides were excluded from fragmentation and dynamic exclusion was set to 60 sec to prevent oversampling of same peptides.

#### Processing of mass spectrometry data and statistical analysis

Data analysis was carried out with MaxQuant v1.6.14.0^73^ using an organism-specific database extracted from Uniprot.org under default settings. Identified false discovery rate (FDR) cutoffs were 0.01 on peptide level and on protein level. The match between runs (MBR) option was enabled to transfer peptide identifications across RAW files based on accurate retention time and m/z. The fractions were set in a way that MBR was only performed within each condition. Label-free quantification (LFQ) was done using a label free quantification approach based on the MaxLFQ algorithm^74^. A minimum of two quantified peptides per protein was required for protein quantification. Adapted from the Perseus recommendations^73^, protein groups with a non-zero LFQ intensity in 70% of the samples of at least one of the conditions were used for statistics. LFQ values were normalized via variance stabilization normalization^75^. Based on the Perseus recommendations, missing LFQ values being completely absent in one condition were imputed with random values drawn from a downshifted (2.2 standard deviation) and narrowed (0.3 standard deviation) intensity distribution of the individual samples. For missing LFQ values with no complete absence in one condition, the R package missForest v1.4 was used for imputation^76^.

### scRNA-seq analysis

#### scRNA-seq library preparation and next-generation sequencing

Sorted cells were counted using the Luna-FL automated cell counter (Logos Biosystems), and 18,000 – 20,000 cells per channel were loaded onto a Chromium controller (10x Genomics), except for GFP^+^ cells, which were loaded completely without counting due to low cell number. scRNA-seq libraries were prepared using Chromium Next GEM Single Cell 3’ v2 (Fig. 1–3 and S1-3) or Chromium Next GEM Single Cell 3’ v3.1 (all other figures) and following the manufacturer’s protocol (10x Genomics). Libraries were analyzed and quantified using TapeStation D1000 screening tapes (Agilent) and Qubit HS DNA quantification kit (Thermo Fisher Scientific) before sequencing with a NextSeq 500 (Illumina) (Fig. 1–3 and S1-3) or NovaSeq 6000 (Illumina) (all other figures). Detailed information for each scRNA-seq run can be found in Supplementary Table 15.

#### Processing of scRNA-seq data

Raw sequencing data was processed with 10x Genomics Cell Ranger v3.1.0^77^. The reads were aligned to the mm10 reference genome v1.2.0 provided by Cell Ranger and, additionally, to a custom reference genome to determine the expression of Scgb1a1, Sftpc, and DTA transgenes in the SRC and SfSRC mice. SoupX v1.5.0^78^ was used to remove ambient RNA contamination. To avoid underestimation of the global contamination fraction, manual gene lists were used for the sequencing runs of GFP^+^ mesenchymal cells on day 2 and 3 (Sftpc, Sftpa1, Sftpb, Sftpd, Dcn, Col1a1, Col1a2, Cldn3, Cldn18, Cldn2, Cldn4) and epithelial cells on day 4 (Sftpc, Sftpa1, Sftpb, Sftpd, Dcn, Scgb3a1, Foxj1) after tamoxifen administration in the SRC model. SoupX was otherwise run with default parameters. Further processing and analysis of count tables was performed with Seurat v3.2.1^79^. Poor quality cells were filtered out based on high content of mitochondrial genes (4 to 7.5%, depending on the sample) and low total number of features (500 to 2500, depending on the sample), before integrating different samples.

#### Sample integration, cell cycle regression, dimensionality reduction, clustering, and doublets exclusion

Individual samples were integrated with Seurat with IntegrateData() using 2,000 anchor features and integrating all common features between samples. Cell cycle phase was calculated by adapting the Seurat function CellCycleScoring() to use the GFP^+^ sorted cells as a reference. Cell cycle scores were then regressed out during data scaling with ScaleData() to mitigate the effects of cell cycle heterogeneity in the datasets from Fig. 3, 4, 7, S3, S4, and S7. UMAP dimensionality reduction and nearest-neighbor graphs were calculated using the top 30 principle components. Cells were then clustered with FindClusters() with a resolution between 0.3 and 1.5, depending on cell number. Cell doublets were identified using scDblFinder v1.2.0^80^, and clusters of cells composed of doublets were excluded from the analysis. Similar clusters were merged, taking into consideration the phylogenetic tree calculated with BuildClusterTree(). Cells were identified based on the expression of known marker genes and, when present, clusters composed of endothelial (Pecam^+^) or hematopoietic (Ptprc^+^) cells were excluded from the analysis. In the case of mesenchymal cell analysis, only clusters expressing Col1a1 were kept.

#### Differential gene expression and gene set enrichment analysis

Differential gene expression between different cell types (markers) was calculated with FindMarkers(test.use=“roc”, only.pos=TRUE, min.pct=0.2), and differentially gene expression between cells from non-injured and injured lungs was calculated with FindMarkers(test.use=“MAST”, min.pct=0.2). Common marker genes of DTA^+^ epithelial cells were calculated separately for downregulated and upregulated genes by intersecting the DEGs previously calculated for each population with FindMarkers(test.use=“MAST”, ident.1=“DTA^+^_cell_type”, ident.2=c(“all populations excluding DTA+ cells”, min.pct=0.2). Only DEGs with a p_adj <0.05 were considered. Heatmaps for DEGs were generated with Seurat’s function DoHeatmap(). GSEA was performed using Metascape v3.5 ^81^ with a p-value cutoff of 10E-6. Activities of 14 pathways were inferred with PROGENy v1.10.0 ^82^ using organism=“Mouse”, scale=FALSE and otherwise default values. Scores were scaled and centered using Seurat’s ScaleData(). Heatmap showing pathway activity for each cell cluster at day 2 was drawn with ComplexHeatmap v2.4.3^83^.

#### Transcriptome correlation between cell types

For every cell type combination of day 2 samples, Spearman’s rank correlation coefficient was calculated from the average normalized expression of all genes. Heatmap was drawn with ComplexHeatmap v2.4.3^83^.

#### Expression of human lung diseases signatures

To infer the expression of human lung diseases signatures in mouse epithelial cells, SRC mouse features were converted to their human orthologs using bioDBnet^84^ and a “humanized” Seurat object was generated using the same parameters as for the mouse object. Single-cell scores were calculated with AddModuleScore() using gene signatures for asthma, non-small cell lung carcinoma, and influenza from DisGeNET v7^48^, and chronic obstructive pulmonary disease^85^, COVID-19 bronchial epithelial cells^51^, and pulmonary fibrosis^86^ from MSigDB v7.5.1^49^’^50^ databases. Expression of the different signatures by each SRC epithelial population at day 0 and day 2 was drawn with the Seurat’s RidgePlot() function.

#### Integration with COVID-19 dataset and DEGs analysis

“Humanized” Seurat objects from SRC mouse epithelial cells at day 0 and day 2 were integrated with lung epithelial cells from controls and COVID-19 patients selected based on the author’s “Epithelial cells” annotation in the “cell_type_main” identity class^4^. FindIntegrationAnchors() was run with a k.filter of 100 due to the low cell number in some patient samples, and SRC mouse samples were used as reference. UMAP dimensional reduction and nearest-neighbor graphs were calculated using the top 30 principle components, and clusters were calculated with FindClusters() and a resolution of 0.5. Cell type identification was done according to the authors’ annotation (“cell_type_fine” identity class) of the majority of the cells in each cluster. When several clusters with the same cell type were identified, a number was added as a suffix. DEGs in DTA^+^ like cells were calculated with FindMarkers(test.use=“MAST, min.pct=0.2, ident.1=“DTA^+^ like cells”) and considering only epithelial cells from COVID-19 patients. Only DEGs with a p_adj <0.05 were considered.

#### Similarity to previously described epithelial cell types

To assess the similarity between our cell populations and mouse lung epithelial cells previously identified, fast gene set enrichment analysis was done as described previously (Laughney *et al*., 2020). Briefly, marker genes for all SfSRC mouse epithelial cell populations using all timepoints were calculated with FindMarkers(test.use=“MAST”, logfc.threshold=-Inf, min.pct=-Inf), and housekeeping genes listed in Laughney *et al*^87^ were excluded. Genes were ranked according to average log FC, and the top 50 genes from previously described epithelial cell types were used to calculate the normalized enrichment score (NES) for each SfSRC mouse epithelial population.

#### Projection of cells from the SRC model onto the SfSRC model

To compare the cell populations present in SRC mice versus SfSRC mice, SingleCellExperiment^88^ objects were generated and SfSRC cells were projected onto SRC cells using scmapCell()^89^. Cell assignment to SRC cell types was done with scmapCell2Cluster() and drawn with getSankey() from scmap v1.10.0 package^89^.

#### Expression of chemokines

Chemokine genes considered for analysis were taken from MGI GO TERM “Chemokine activity”. scRNA-seq data to assess chemokine expression in homeostasis by lung cell types, including mesenchymal, immune, and endothelial cells, was taken from the Mouse Cell Atlas (Fig. S5b)^53^.

#### Cell-cell communication analysis

Intercellular interactions were inferred in mesenchymal (Fig. 2 cells) and epithelial cells (Fig. 4 cells) of the SRC model before (day 0) and after tamoxifen administration (day 2) with CellChat v1.1.0^35^ following the official workflow and using standard parameters. The analysis was based on expression of ligand-receptor pairs from the CellChat mouse database, which we manually adjusted to exclude ligand-receptor pairs not supported by literature and to include interactions playing a role in intercellular junctions and mesenchymal cells of the lung (adjusted database is provided in Supplementary Table 14). Neuroendocrine cells were excluded from the analysis due to low cell numbers. For each timepoint, preprocessing of normalized count tables was performed with identifyOverExpressedGenes(), identifyOverExpressedInteractions() and projectData(). The cell-cell communication network was inferred using computeCommunProb(), which by default requires 25% of the cells per group to express the ligand or receptor gene. Summarizing analyses were performed with computeCommunProbPathway(), aggregateNet() and netAnalysis_computeCentrality(). The summed incoming and outgoing interactions strengths were obtained with netAnalysis_signalingRole_scatter() and scaled to the maximum summed interaction strength at the respective time point. For comparison of the signaling pathways between the two timepoints, the CellChat objects were merged with liftCellChat() and mergeCellChat().

#### Force-directed graph and PAGA analysis

Connectivities between different cell types were analyzed with the Scanpy package^90^. PCA and a neighborhood graph were computed with pp.pca() and sc.pp.neighbors() using n_neighbors=10 and n_pcs=20. The force-directed graph was drawn with sc.tl.draw_graph(). For the PAGA analysis, the data was restricted to non-dividing cell clusters, on which the PCA and neighborhood graph were re-calculated using the same functions and parameters as above. Connectivities were quantified with tl.paga()^91^.

#### Diffusion maps

Diffusion maps restricted to cell type subsets of interest were calculated with Scanpy^90^. The data was restricted to the cell type subset and the top 2,000 variable genes with a minimum count of 10. To avoid obtaining cell cycle-related genes, dividing cell clusters and cells in the G2/M or S phase were removed for the calculation of the top variable genes. PCA and the neighborhood graph were calculated with pp.pca() and sc.pp.neighbors() using n_neighbors=15 and n_pcs=20. The diffusion map was calculated with tl.diffmap().

#### RNA velocity

Spliced and unspliced read counts per gene were obtained with Velocyto v0.17.17^92^ and merged with the pre-processed normalized and log-transformed count data from Seurat. To reduce the influence of possible variable kinetic rates between different cell types or states^93^, RNA velocities were calculated separately on a reduced data set containing only cell types along the transition pathway using scVelo v0.2.2^94^. First and second order moments were calculated using scvelo.pp.moments() with default parameters, velocities were calculated with scvelo.tl.velocity() in “dynamical” mode, and a velocity graph was constructed with scvelo.tl.velocity_graph(). The velocities were projected and visualized on diffusion map embeddings (see above) using scvelo.pl.velocity_embedding_grid(density = 0.5).

#### Trajectory inference and differential expression analysis

In the diffusion maps, trajectory analysis was performed with Monocle 3 v0.2.2^95^ using default parameters, unless otherwise specified. The scale of the diffusion map DC values was adjusted by multiplying by 100. Cells in the diffusion map were clustered using cluster_cells() with parameters partition_qval=0.05 and num_iter=1. Trajectories were inferred using learn_graph(). Cells were assigned a pseudotime with the order_cells() function, for which root cells were manually chosen according to prior knowledge. The differential expression analysis was manually restricted to cells along the respective branch of interest and carried out using the function graph_test() with the parameter neighbor_graph=“principal_graph”. The expression was then scaled to the cell subset and smoothed using the loess() function in R with span set to 0.5. Heatmaps of the top 1,000 DEGs (sorted by q-value and Morans’s I) were drawn with ComplexHeatmap v2.4.3^83^. Mouse transcription factors in the DEGs were annotated with AnimalTFDBv3.0^96^.

## Acknowledgements

We thank the Heidelberg University Nikon Imaging Center, the DKFZ Transgenic Core Facility, the DKFZ Center for Preclinical Research, the DKFZ Light Microscopy Core Facility, the DKFZ Flow Cytometry Core Facility, the DKFZ Genomics and Proteomics Core Facility, the EMBL Genomics Core Facility, and the Single-cell Open Lab for valuable technical assistance and discussions. We thank Roman Kurley, Jana Kress, Gelsomina Kaufhold, Dorothee Terhardt, Alexandra Buse, and Lea Jopp-Saile for technical assistance, and Rocio Sotillo for providing the SPCcreT2rtTA mice. This study was supported by the German Research Foundation (DFG) grant SFB873 to L.R.M., C.S., and H.G.. L.V. acknowledges support of the Spanish Ministry of Science and Innovation to the EMBL partnership, the Centro de Excelencia Severo Ochoa and the CERCA Programme/Generalitat de Catalunya.

## Author contributions

Cl.S. and H.G. designed the study; Cl.S. supervised the work; L.V. provided expertise for and cosupervised data analysis; L.R.M. performed most experiments and computational analyses; M.M. performed experiments and computational analyses related to mesenchymal cells; L.S. aligned reads and generated feature-barcode matrices, performed computational analyses related to cellcell interaction, chemokine expression, and differentiation trajectories; S.T. performed preliminary computational analyses related to differentiation trajectories; Ch.S. pre-processed previously published datasets and curated the data used to evaluate cell-cell interaction; C.E. provided computational support for the scRNA-seq data analysis; S.F. contributed valuable resources; L.R.M., M.M., L.S., and Cl.S. wrote the manuscript with input from other authors.

## Additional information

Supplemental figures and tables are provided with this manuscript.

