## Supplementary figures and table legends for "Cell division tracing combined with single-cell transcriptomics reveals new cell types and differentiation paths in the regenerating mouse lung"

### SUPPLEMENTARY MATERIAL

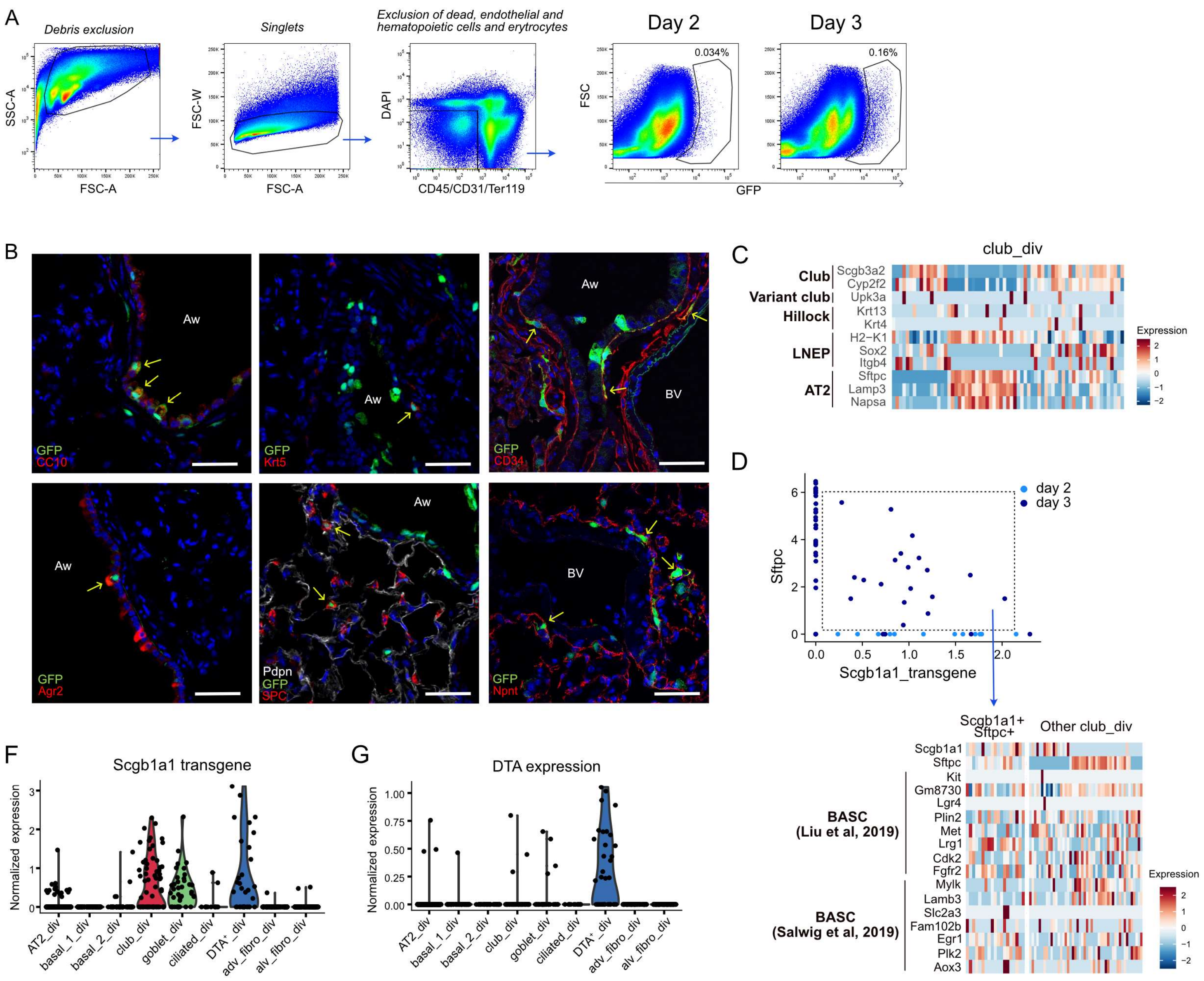


**Fig. S1: Characterization of dividing cells after targeted depletion of *Scgb1a1*⁺ cells.**

**A.** Gating strategy used to sort GFP⁺ cells. **B.** IF staining of dividing GFP⁺ cells (green) combined with markers for club cells (CC10, red), goblet cells (Agr2, red), basal cells (Krt5, red), AT1 cells (Pdpn, white), AT2 cells (SPC, red), adventitial fibroblasts (CD34, red), and alveolar fibroblasts (Npnt, red). Nuclei are stained with DAPI (blue). All images were taken at day 3 after tamoxifen injection. Aw: airway; BV: blood vessel. Arrows are pointing to GFP+ cells co-stained with cell type specific markers. Scale bar: 50 µm. **C.** Heatmap of the main marker genes for previously described progenitor club cell types, club cells, and AT2 cells in club_div cells at day 2 and 3. **D**. Upper panel: Co-expression of the Scgb1a1 transgene and *Sftpc* in club_div cells. Lower panel: Expression of bronchioalveolar stem cell (BASC) marker genes in club_div cells co-expressing the Scgb1a1_transgene and *Sftpc* and in other club_div cells. **F, G.** Violin plots showing the expression of the Scgb1a1 transgene **(F)** and DTA **(G)** in the dividing populations at day 2 and 3.

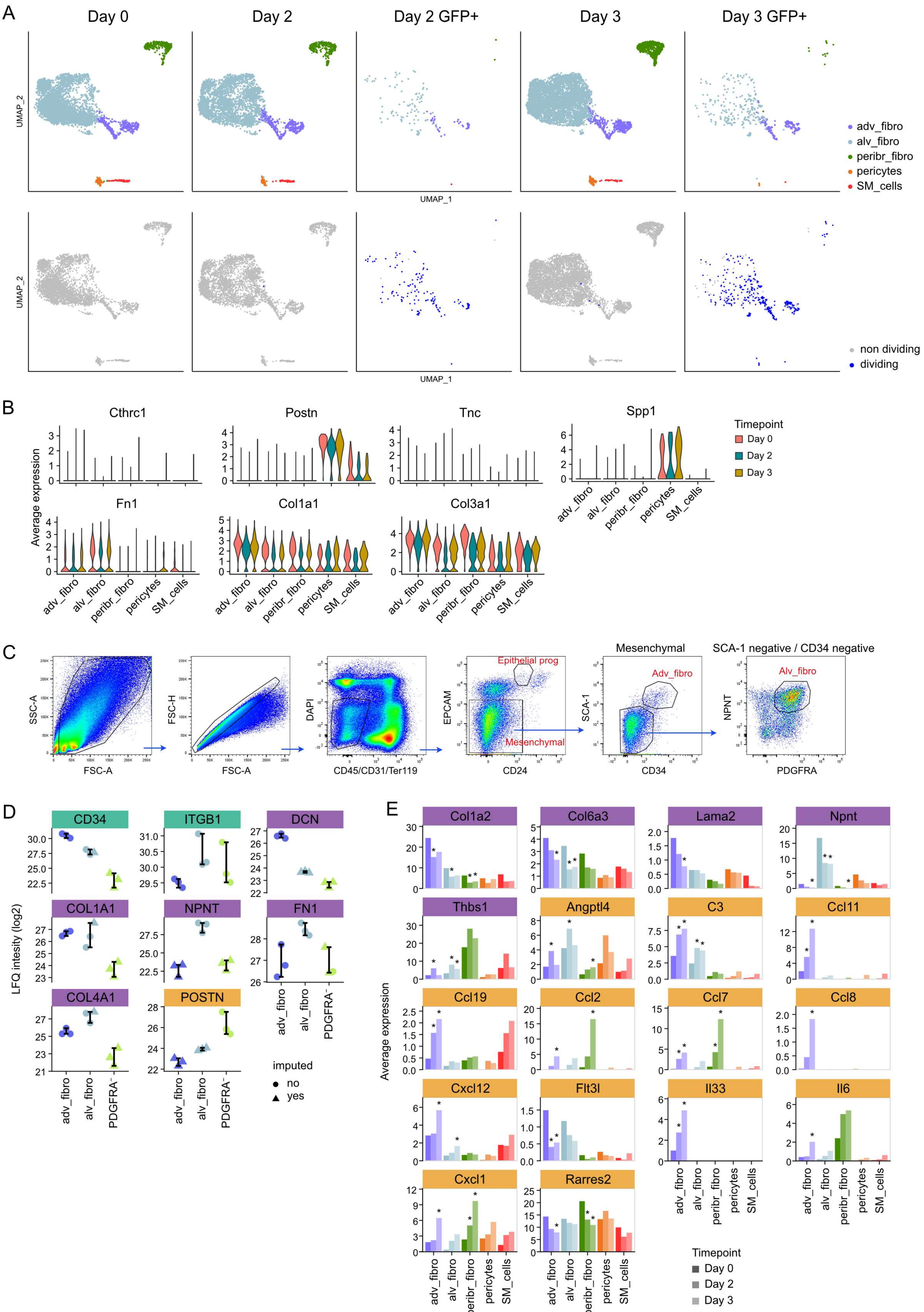
A
Day 0
Day 2
Day 2 GFP+
Day 3
Day 3 GFP+
UMAP_2
UMAP_1
adv_fibro
alv_fibro
peribr_fibro
pericytes
SM_cells
non dividing
dividing
B
Cthrc1
Postn
Tnc
Spp1
Fn1
Col1a1
Col3a1
Average expression
Timepoint
Day 0
Day 2
Day 3
C
SSC-A
FSC-A
FSC-H
DAPI
CD45/CD31/Ter119
EPCAM
CD24
Epithelial prog
Mesenchymal
SCA-1
CD34
Adv_fibro
SCA-1 negative / CD34 negative
NPNT
PDGFRA
Alv_fibro
D
CD34
ITGB1
DCN
COL1A1
NPNT
FN1
COL4A1
POSTN
LFQ intesity (log2)
PDGFRA⁻
imputed
no
yes
E
Col1a2
Col6a3
Lama2
Npnt
Thbs1
Angptl4
C3
Ccl11
Ccl19
Ccl2
Ccl7
Ccl8
Cxcl12
Flt3l
Il33
Il6
Cxcl1
Rarres2
Average expression
Timepoint
Day 0
Day 2
Day 3

**Fig. S2: Characterization of mesenchymal cell activation after epithelial injury in SRC mice.**

**A.** UMAP embedding of mesenchymal cells and dividing $GFP^+$ mesenchymal cells on the indicated days. **B.** Violin plots showing the expression of fibrosis-associated genes in the different mesenchymal populations on day 0, 2, and 3. **C.** Gating strategy used to sort epithelial progenitor cells (epithelial prog), adventitial fibroblasts (adv_fibro), and alveolar fibroblasts (alv_fibro). **D.** Protein expression in log2 label-free quantification (LFQ) intensity of upregulated ligands in adventitial fibroblasts, alveolar fibroblasts, and PDGFRA negative cells (see Fig. 2E) at day 0 for which protein data were available. **E.** Average normalized expression of all ligands in the modified CellChat ligand-receptor database that are differentially expressed between day 0 and day 2 or day 3 in adv_fibro. *Bmp4*, *Tgfb3,* and *Mif* (included in Fig. 2F) were excluded from this figure. Asterisks denote differential expression on day 2 or 3 (FC >=1.5, p_adj <=0.05). **D, E.** Ligand genes are sorted by the possible type of interaction (yellow: secreted signaling, purple: ECM-receptor, green: cell-cell contact). **A, B, E.** scRNA-seq analysis was done using total mesenchymal cells on day 0 (n=2 mice), day 2 (n=3 mice), and day 3 (n=3 mice), and $GFP^+$ mesenchymal cells on day 2 (n=12 mice) and day 3 (n=3 mice).

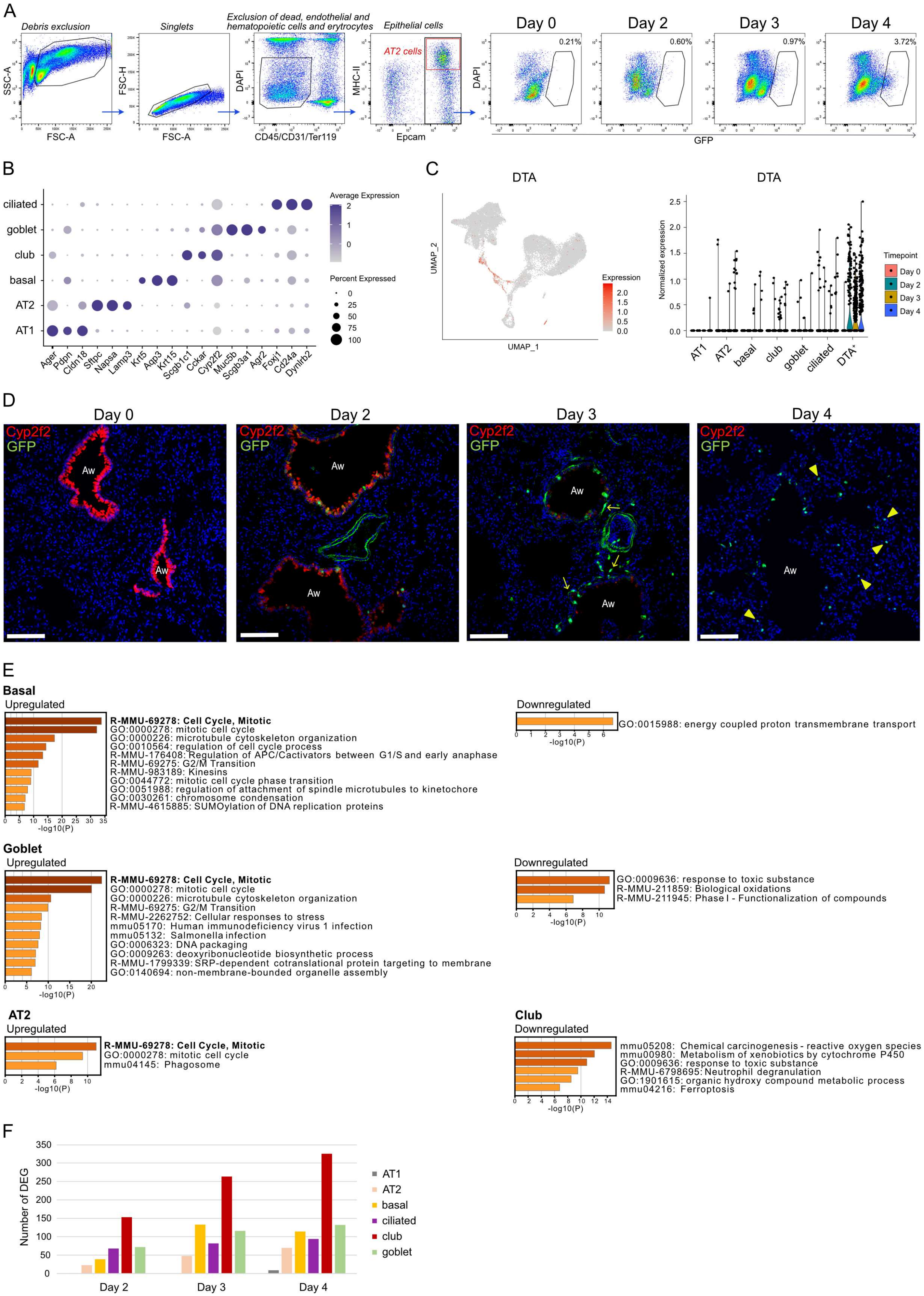
A
Debris exclusion
Singlets
Exclusion of dead, endothelial and hematopoietic cells and erytrocytes
Epithelial cells
AT2 cells
Day 0
Day 2
Day 3
Day 4
0.21%
0.60%
0.97%
3.72%
SSC-A
FSC-A
FSC-H
FSC-A
DAPI
CD45/CD31/Ter119
MHC-II
Epcam
DAPI
GFP
B
ciliated
goblet
club
basal
AT2
AT1
Ager
Pdpn
Cldn18
Sftpc
Napsa
Lamp3
Krt5
Aqp3
Krt15
Scgb1c1
Cckar
Cyp2f2
Muc5b
Scgb3a1
Agr2
Foxj1
Cd24a
Dynlrb2
Average Expression
2
1
0
Percent Expressed
0
25
50
75
100
C
DTA
UMAP_2
UMAP_1
Expression
2.0
1.5
1.0
0.5
0.0
DTA
Normalized expression
2.5
2.0
1.5
1.0
0.5
0.0
AT1
AT2
basal
club
goblet
ciliated
DTA+
Timepoint
Day 0
Day 2
Day 3
Day 4
D
Day 0
Day 2
Day 3
Day 4
Cyp2f2
GFP
Aw
E
Basal
Upregulated
R-MMU-69278: Cell Cycle, Mitotic
GO:0000278: mitotic cell cycle
GO:0000226: microtubule cytoskeleton organization
GO:0010564: regulation of cell cycle process
R-MMU-176408: Regulation of APC/Cactivators between G1/S and early anaphase
R-MMU-69275: G2/M Transition
R-MMU-983189: Kinesins
GO:0044772: mitotic cell cycle phase transition
GO:0051988: regulation of attachment of spindle microtubules to kinetochore
GO:0030261: chromosome condensation
R-MMU-4615885: SUMOylation of DNA replication proteins
-log10(P)
Downregulated
GO:0015988: energy coupled proton transmembrane transport
-log10(P)
Goblet
Upregulated
R-MMU-69278: Cell Cycle, Mitotic
GO:0000278: mitotic cell cycle
GO:0000226: microtubule cytoskeleton organization
R-MMU-69275: G2/M Transition
R-MMU-2262752: Cellular responses to stress
mmu05170: Human immunodeficiency virus 1 infection
mmu05132: Salmonella infection
GO:0006323: DNA packaging
GO:0009263: deoxyribonucleotide biosynthetic process
R-MMU-1799339: SRP-dependent cotranslational protein targeting to membrane
GO:0140694: non-membrane-bounded organelle assembly
-log10(P)
Downregulated
GO:0009636: response to toxic substance
R-MMU-211859: Biological oxidations
R-MMU-211945: Phase I - Functionalization of compounds
-log10(P)
AT2
Upregulated
R-MMU-69278: Cell Cycle, Mitotic
GO:0000278: mitotic cell cycle
mmu04145: Phagosome
-log10(P)
Club
Downregulated
mmu05208: Chemical carcinogenesis - reactive oxygen species
mmu00980: Metabolism of xenobiotics by cytochrome P450
GO:0009636: response to toxic substance
R-MMU-6798695: Neutrophil degranulation
GO:1901615: organic hydroxy compound metabolic process
mmu04216: Ferroptosis
-log10(P)
F
Number of DEG
350
300
250
200
150
100
50
0
Day 2
Day 3
Day 4
AT1
AT2
basal
ciliated
club
goblet

**Fig. S3: Transcriptional changes in epithelial lung cells after *Scgb1a1*+ cell depletion.**

**A.** Gating strategy used to sort epithelial cells, while keeping only 10% of AT2 cells (MHC II+) in relation to the total number of cells (red gate), and gating strategy to sort GFP+ epithelial cells. Percentage of GFP+ epithelial cells in relation to total epithelial cells on day 0 to 4 is indicated. **B.** Dot plot showing the average scaled expression of cell type-associated marker genes across the epithelial populations. **C.** UMAP embedding (left) and violin plot (right) showing the expression of DTA in epithelial cell types from day 0 to day 4. **D.** IF staining of SRC mouse lungs with Cyp2f2 (red; club cells), CycB1-GFP (green; dividing cells), and DAPI (blue; nuclei) before (day 0) and 2, 3, and 4 days after injecting tamoxifen on two consecutive days. GFP+ spindle-like cells (arrows) and GFP+ alveolar cells (arrow heads) can be observed from day 3. Aw: airway. Scale bar: 100 µm. Images are representative of at least three animals analyzed per group. **E.** GSEA of DEGs on day 2, 3, and 4 compared to day 0 (FC >=2, p_adj <0.05) in the different epithelial populations. **F.** Number of DEGs at the indicated days as compared to day 0 for all epithelial cell types (FC >2, p_adj <0.05).

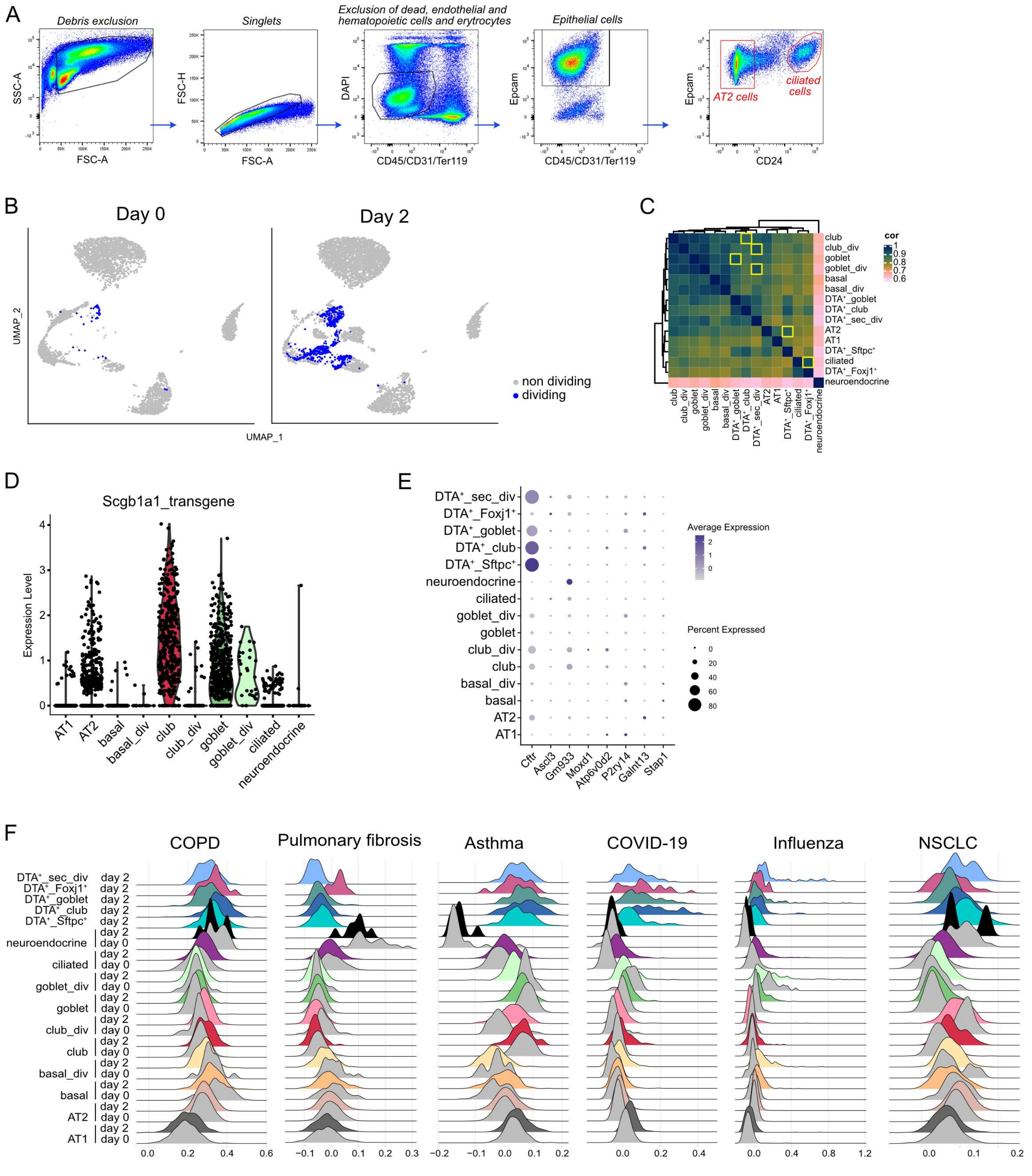


**Fig. S4: Characterization of DTA⁺ epithelial cells and comparison to transcriptional profiles of human lung diseases.**

**A.** Gating strategy used to sort epithelial cells, while keeping only 10% of AT2 cells (CD24$^{-}$) and 10% of ciliated cells (CD24$^{high}$) in relation to the total number of cells (red gates). **B.** UMAP embedding showing the distribution of dividing and non-dividing cells. **C.** Heatmap of the transcriptome Sperman correlation coefficient (cor) between all epithelial cell types on day 2. Yellow boxes highlight the closest population to each DTA$^{+}$ cell type. **D.** Violin plot of the Scgb1a1 transgene expression on day 0 across all epithelial cells in SRC mice. **E.** Dot plot of ionocyte marker gene expression in all epithelial cell populations at day 2. **F.** Ridge plots showing the expression of human lung diseases signatures in SRC mouse epithelial cells on day 0 and day 2.

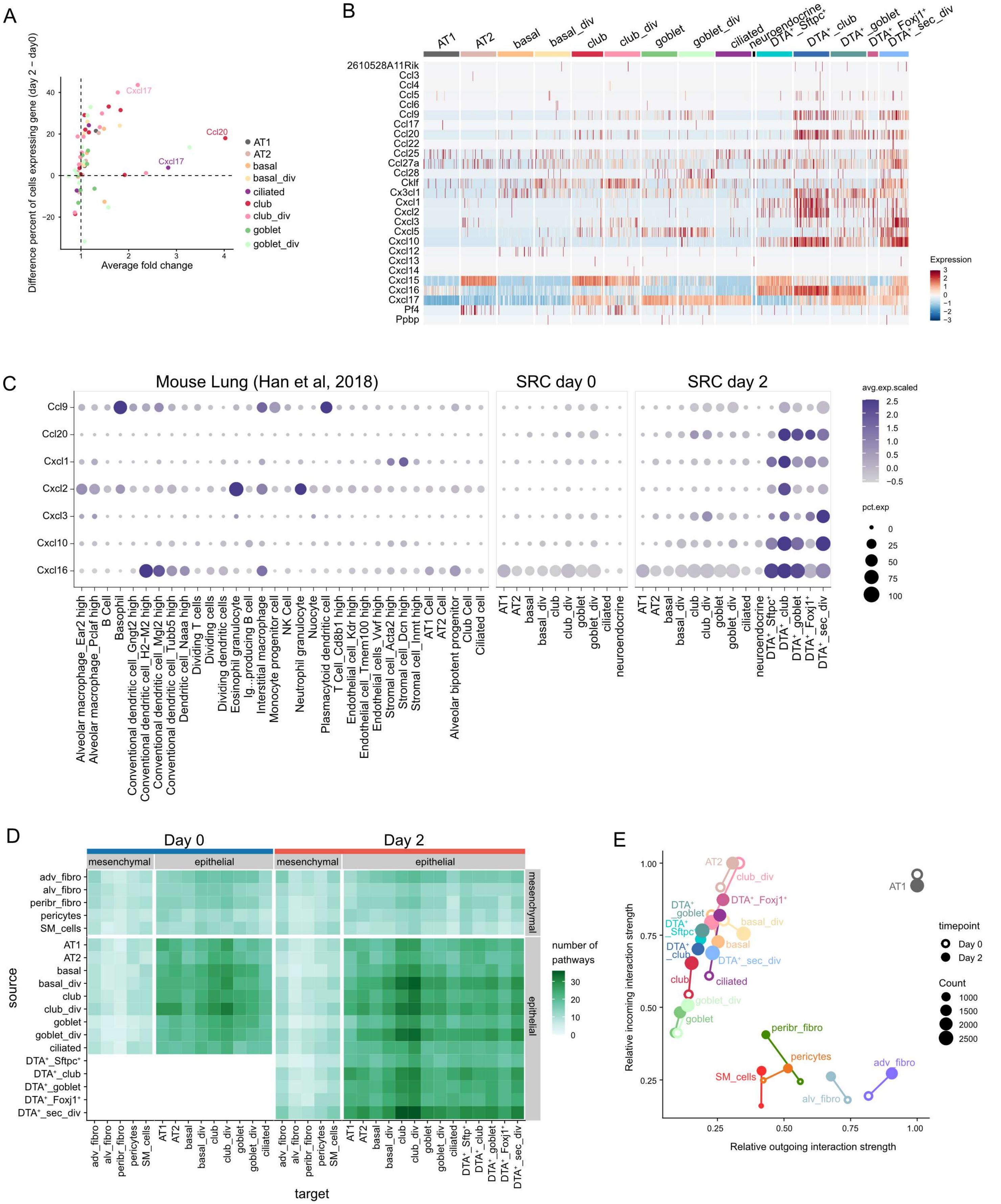
A
Difference percent of cells expressing gene (day 2 − day0)
Average fold change
Cxcl17
Ccl20
Cxcl17
AT1
AT2
basal
basal_div
ciliated
club
club_div
goblet
goblet_div
B
AT1
AT2
basal
basal_div
club
club_div
goblet
goblet_div
ciliated
neuroendocrine
DTA⁺_Sftpc⁺
DTA⁺_club
DTA⁺_goblet
DTA⁺_Foxj1⁺
DTA⁺_sec_div
2610528A11Rik
Ccl3
Ccl4
Ccl5
Ccl6
Ccl9
Ccl17
Ccl20
Ccl22
Ccl25
Ccl27a
Ccl28
Cklf
Cx3cl1
Cxcl1
Cxcl2
Cxcl3
Cxcl5
Cxcl10
Cxcl12
Cxcl13
Cxcl14
Cxcl15
Cxcl16
Cxcl17
Pf4
Ppbp
Expression
C
Mouse Lung (Han et al, 2018)
SRC day 0
SRC day 2
Ccl9
Ccl20
Cxcl1
Cxcl2
Cxcl3
Cxcl10
Cxcl16
avg.exp.scaled
pct.exp
Alveolar macrophage_Ear2 high
Alveolar macrophage_Pclaf high
B Cell
Basophil
Conventional dendritic cell_Gngt2 high
Conventional dendritic cell_H2−M2 high
Conventional dendritic cell_Mgl2 high
Conventional dendritic cell_Tubb5 high
Dendritic cell_Naaa high
Dividing T cells
Dividing cells
Dividing dendritic cells
Eosinophil granulocyte
Ig...producing B cell
Interstitial macrophage
Monocyte progenitor cell
NK Cell
Neutrophil granulocyte
Nuocyte
Plasmacytoid dendritic cell
T Cell_Cd8b1 high
Endothelial cell_Kdr high
Endothelial cell_Tmem100 high
Endothelial cells_Vwf high
Stromal cell_Acta2 high
Stromal cell_Dcn high
Stromal cell_Inmt high
AT1 Cell
AT2 Cell
Alveolar bipotent progenitor
Club Cell
Ciliated cell
D
Day 0
Day 2
mesenchymal
epithelial
adv_fibro
alv_fibro
peribr_fibro
pericytes
SM_cells
AT1
AT2
basal
basal_div
club
club_div
goblet
goblet_div
ciliated
DTA⁺_Sftpc⁺
DTA⁺_club
DTA⁺_goblet
DTA⁺_Foxj1⁺
DTA⁺_sec_div
source
target
number of pathways
E
Relative incoming interaction strength
Relative outgoing interaction strength
timepoint
Day 0
Day 2
Count
1000
1500
2000
2500

F

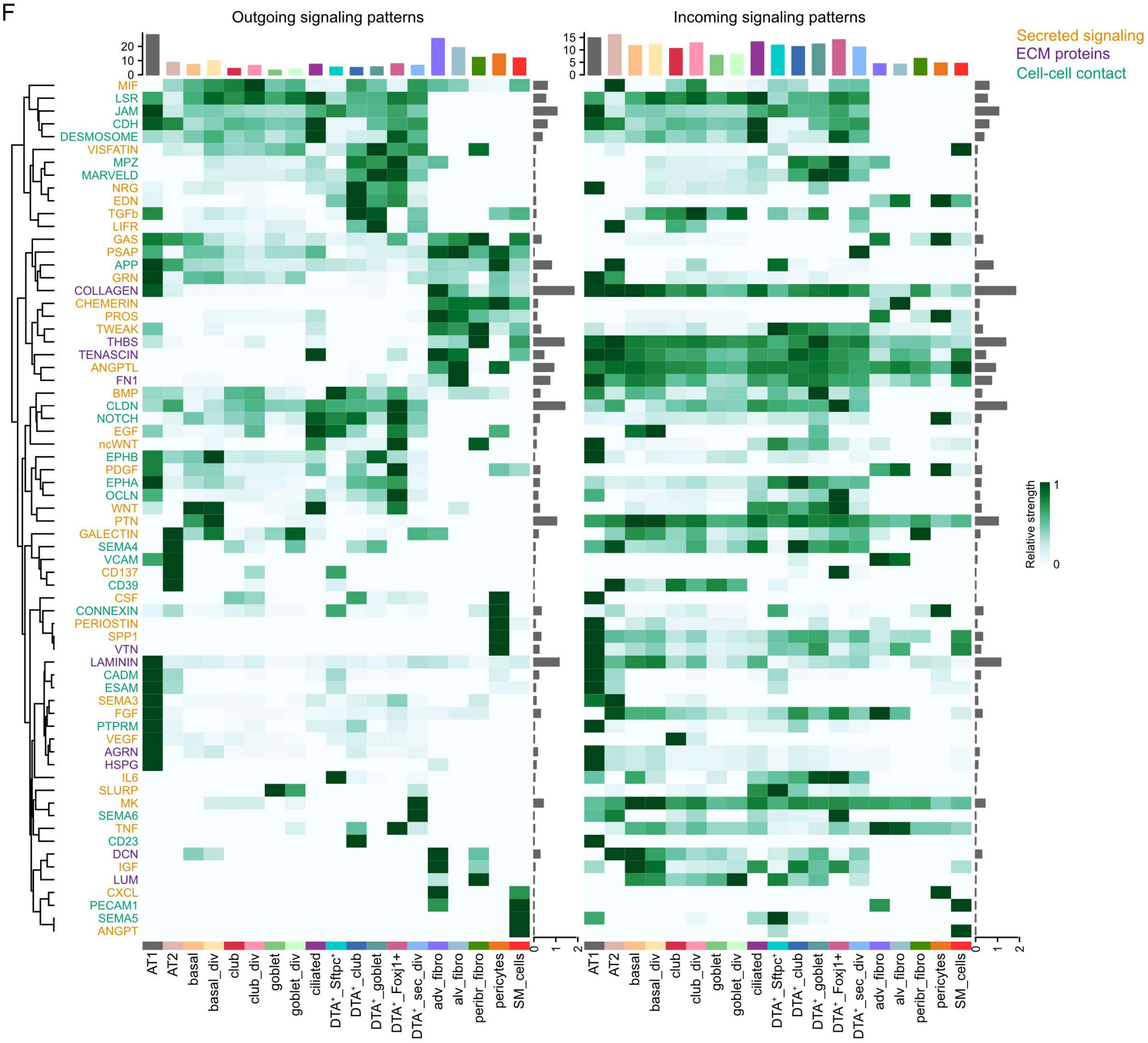

Outgoing signaling patterns
Incoming signaling patterns
Secreted signaling
ECM proteins
Cell-cell contact
Relative strength
MIF
LSR
JAM
CDH
DESMOSOME
VISFATIN
MPZ
MARVELD
NRG
EDN
TGFb
LIFR
GAS
PSAP
APP
GRN
COLLAGEN
CHEMERIN
PROS
TWEAK
THBS
TENASCIN
ANGPTL
FN1
BMP
CLDN
NOTCH
EGF
ncWNT
EPHB
PDGF
EPHA
OCLN
WNT
PTN
GALECTIN
SEMA4
VCAM
CD137
CD39
CSF
CONNEXIN
PERIOSTIN
SPP1
VTN
LAMININ
CADM
ESAM
SEMA3
FGF
PTPRM
VEGF
AGRN
HSPG
IL6
SLURP
MK
SEMA6
TNF
CD23
DCN
IGF
LUM
CXCL
PECAM1
SEMA5
ANGPT
AT1
AT2
basal
basal_div
club
club_div
goblet
goblet_div
ciliated
DTA⁺_Sftpc⁺
DTA⁺_club
DTA⁺_goblet
DTA⁺_Foxj1+
DTA⁺_sec_div
adv_fibro
alv_fibro
peribr_fibro
pericytes
SM_cells


G

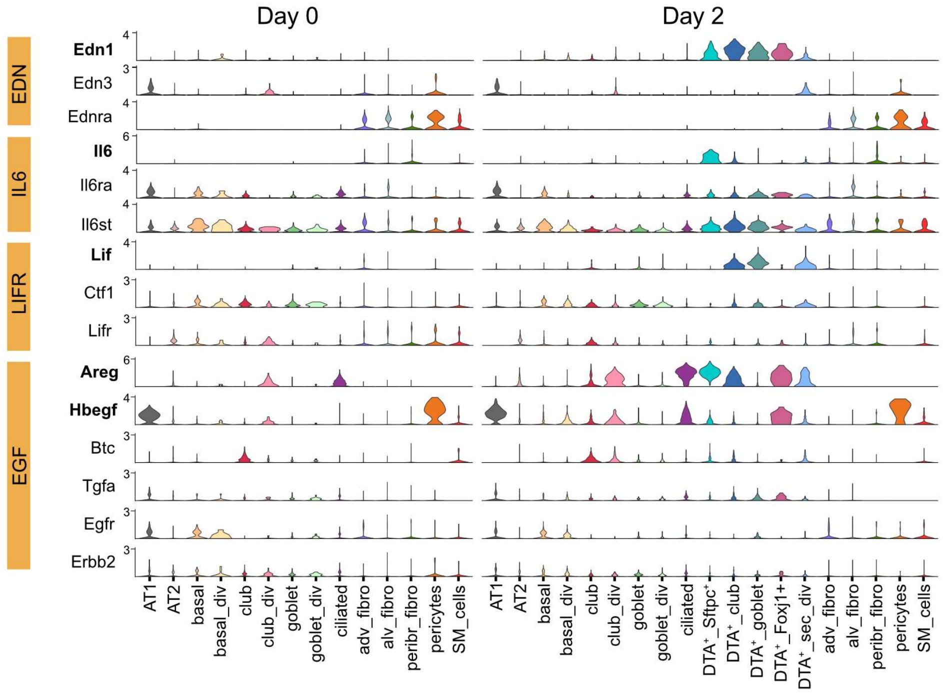

Day 0
Day 2
EDN
Edn1
Edn3
Ednra
IL6
Il6
Il6ra
Il6st
LIFR
Lif
Ctf1
Lifr
EGF
Areg
Hbegf
Btc
Tgfa
Egfr
Erbb2
AT1
AT2
basal
basal_div
club
club_div
goblet
goblet_div
ciliated
adv_fibro
alv_fibro
peribr_fibro
pericytes
SM_cells
DTA⁺_Sftpc⁺
DTA⁺_club
DTA⁺_goblet
DTA⁺_Foxj1+
DTA⁺_sec_div

**Fig. S5: Chemokine expression and cellular crosstalk in the SRC mouse model.**

**A.** Differential expression of chemokines in epithelial lung cells upon injury. Only genes detected in at least 20% of cells in a population in either of the timepoints are shown. Genes with FC > 2 and p_adj < 0.05 are labeled. **B.** Heatmap of chemokine expression in epithelial lung cells on day 2. Scaling of expression was done after downsampling to 100 cells per cell type. Chemokines without counts in any of the cell types were excluded. **C.** Expression of selected chemokines (see Fig. 5B) in lung cells in homeostasis from the mouse atlas of Han *et al.* [53] and in epithelial lung cells of the SRC model on day 0 and day 2. **D.** Number of signaling pathways between any two cell types before (day 0) and after tamoxifen administration (day 2) of the modified CellChat ligand-receptor database. **E.** Summed incoming and outgoing interaction strength of each cell type. The interaction strength was scaled to the maximum summed interaction strength at the respective time point. Circle size is proportional to the number of significant ligand-receptor pairs of the respective cell type at that time point. **F.** Heatmaps of outgoing (left) and incoming (right) signaling patterns per cell type on day 2. The signaling strengths are scaled to the maximum of each row. Column annotations show the summed absolute signaling strengths of each cell type. Row annotations show the log information flow (i.e. summed signaling strength) of each pathway. Signaling pathways are colored by the type of interaction (yellow: secreted signaling, purple: ECM proteins, green: cell-cell contact) and sorted according to the dendrogram of outgoing signaling patterns. **G.** Normalized expression of the main ligand and receptor genes contributing to the EDN, IL6, LIFR, and EGF signaling pathways on day 0 and day2.

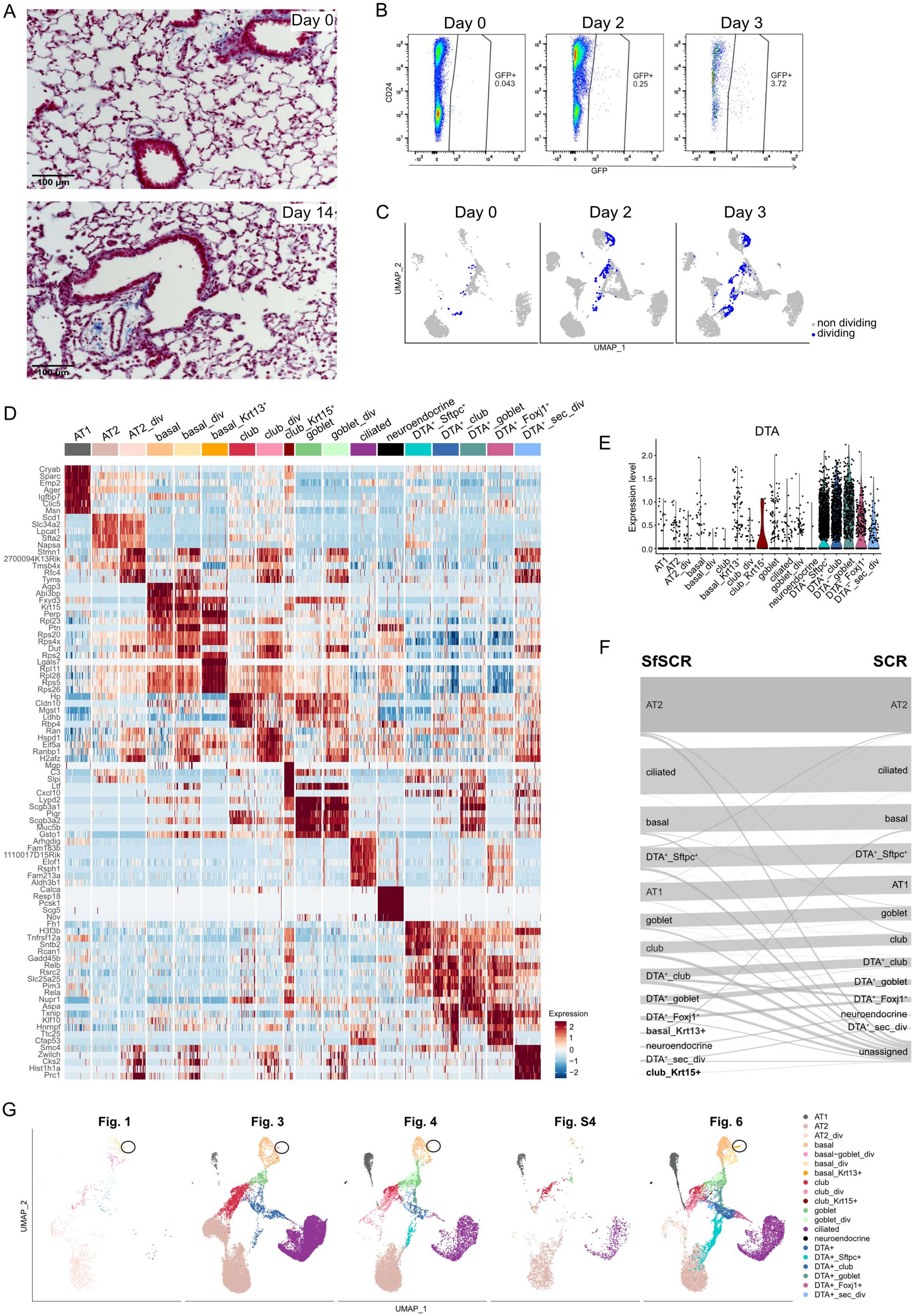
A
Day 0
100 μm
Day 14
100 μm
B
Day 0
Day 2
Day 3
CD24
GFP+
0.043
GFP+
0.25
GFP+
3.72
GFP
C
Day 0
Day 2
Day 3
UMAP_2
UMAP_1
non dividing
dividing
D
AT1
AT2
AT2_div
basal
basal_div
basal_Krt13+
club
club_div
club_Krt15+
goblet
goblet_div
ciliated
neuroendocrine
DTA+_Sftpc+
DTA+_club
DTA+_goblet
DTA+_Foxj1+
DTA+_sec_div
Cryab
Sparc
Emp2
Ager
Igfbp7
Clic5
Msn
Scd1
Slc34a2
Lpcat1
Sfta2
Napsa
Stmn1
2700094K13Rik
Tmsb4x
Rfc4
Tyms
Aqp3
Abi3bp
Fxyd3
Krt15
Perp
Rpl23
Ptn
Rps20
Rps4x
Dut
Rps2
Lgals7
Rpl11
Rpl28
Rps5
Rps26
Hp
Cldn10
Mgst1
Ldhb
Rbp4
Ran
Hspd1
Eif5a
Ranbp1
H2afz
Mgp
C3
Slpi
Ltf
Cxcl10
Lypd2
Scgb3a1
Pigr
Scgb3a2
Muc5b
Gsto1
Arhgdig
Fam183b
1110017D15Rik
Elof1
Rsph1
Fam213a
Aldh3b1
Calca
Resp18
Pcsk1
Scg5
Nov
Fh1
H3f3b
Tnfrsf12a
Sntb2
Rcan1
Gadd45b
Relb
Rsrc2
Slc25a25
Pim3
Rela
Nupr1
Aspa
Txnip
Klf10
Hnrnpf
Ttc25
Cfap53
Smc4
Zwilch
Cks2
Hist1h1a
Prc1
Expression
2
1
0
−1
−2
E
DTA
Expression level
2.0
1.5
1.0
0.5
0.0
F
SfSCR
SCR
AT2
ciliated
basal
DTA+_Sftpc+
AT1
goblet
club
DTA+_club
DTA+_goblet
DTA+_Foxj1+
basal_Krt13+
neuroendocrine
DTA+_sec_div
club_Krt15+
DTA+_sec_div
unassigned
G
Fig. 1
Fig. 3
Fig. 4
Fig. S4
Fig. 6
UMAP_2
UMAP_1
AT1
AT2
AT2_div
basal
basal−goblet_div
basal_div
basal_Krt13+
club
club_div
club_Krt15+
goblet
goblet_div
ciliated
neuroendocrine
DTA+
DTA+_Sftpc+
DTA+_club
DTA+_goblet
DTA+_Foxj1+
DTA+_sec_div

**Fig. S6: Characterization of lung epithelial cells in the SfSRC mouse model.**

**A.** Masson staining of SfSRC mouse lungs before (day 0) and 14 days after two consecutive tamoxifen injections. Images are representative of at least 3 animals per group. **B.** Percentage of $GFP^+$ epithelial cells in relation to total epithelial cells before (day 0) and two and three days after tamoxifen injection in SfSRC mouse lungs. **C.** UMAP embedding showing dividing and non-dividing cells on day 0, 2, and 3. **D.** Heatmap of the top five marker genes ranked by power across all epithelial populations (all timepoints merged). **E.** Violin plot of DTA expression (all timepoints merged). **F.** Sankey diagram of SfRC mouse epithelial cells projected onto SRC mouse epithelial populations. **G.** UMAP embedding of all scRNA-seq datasets used in this study. Circles depict *Krt13*+ basal cells.

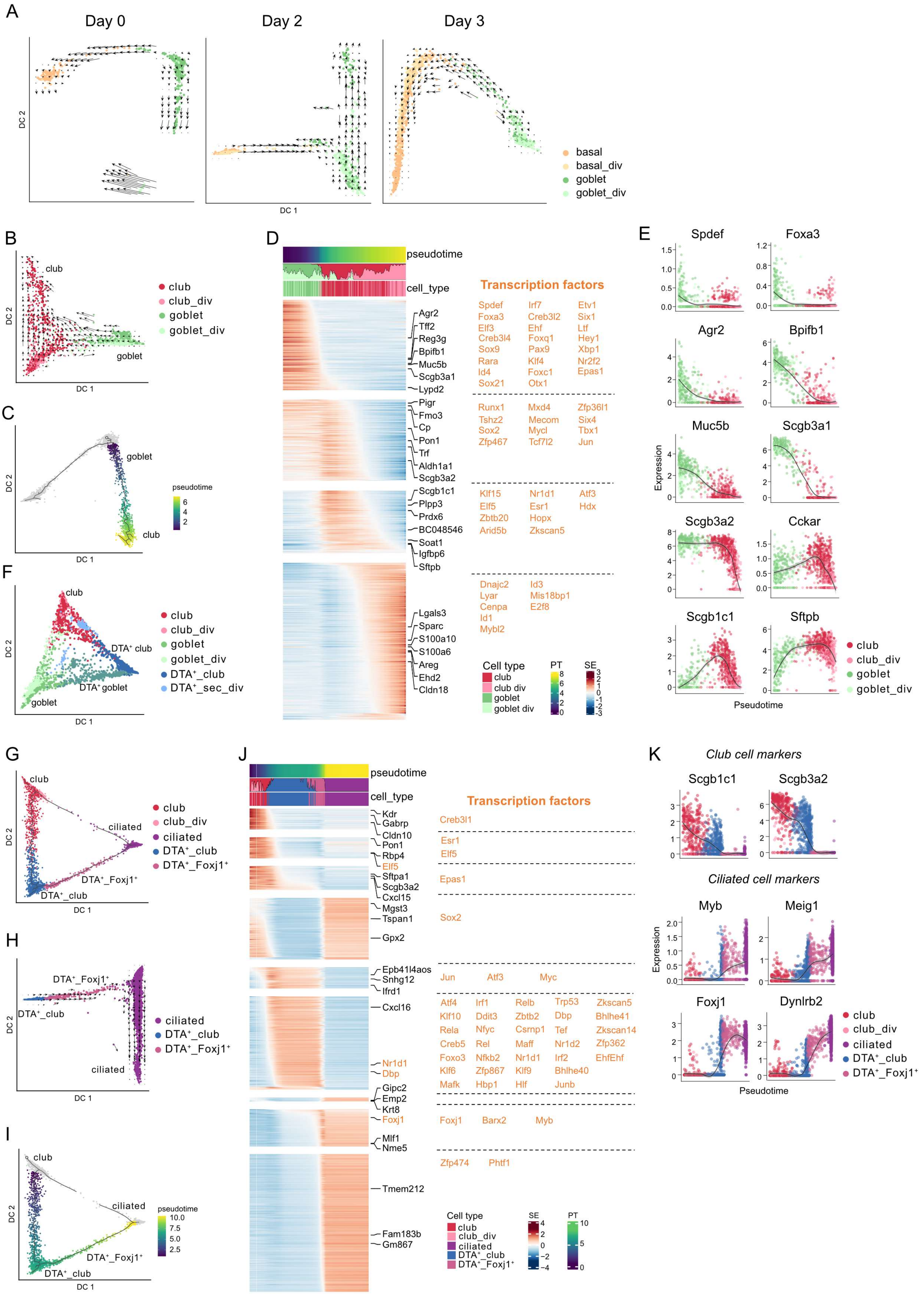
A
Day 0
Day 2
Day 3
DC 2
DC 1
basal
basal_div
goblet
goblet_div
B
club
goblet
club_div
goblet_div
C
pseudotime
D
pseudotime
cell_type
Agr2
Tff2
Reg3g
Bpifb1
Muc5b
Scgb3a1
Lypd2
Pigr
Fmo3
Cp
Pon1
Trf
Aldh1a1
Scgb3a2
Scgb1c1
Plpp3
Prdx6
BC048546
Soat1
Igfbp6
Sftpb
Lgals3
Sparc
S100a10
S100a6
Areg
Ehd2
Cldn18
Transcription factors
Spdef Irf7 Etv1
Foxa3 Creb3l2 Six1
Elf3 Ehf Ltf
Creb3l4 Foxq1 Hey1
Sox9 Pax9 Xbp1
Rara Klf4 Nr2f2
Id4 Foxc1 Epas1
Sox21 Otx1
Runx1 Mxd4 Zfp36l1
Tshz2 Mecom Six4
Sox2 Mycl Tbx1
Zfp467 Tcf7l2 Jun
Klf15 Nr1d1 Atf3
Elf5 Esr1 Hdx
Zbtb20 Hopx
Arid5b Zkscan5
Dnajc2 Id3
Lyar Mis18bp1
Cenpa E2f8
Id1
Mybl2
Cell type
club
club div
goblet
goblet div
PT
SE
E
Spdef
Foxa3
Agr2
Bpifb1
Muc5b
Scgb3a1
Scgb3a2
Cckar
Scgb1c1
Sftpb
Expression
Pseudotime
F
DTA⁺ club
DTA⁺ goblet
DTA⁺_club
DTA⁺_sec_div
G
ciliated
DTA⁺_Foxj1⁺
H
I
J
Kdr
Gabrp
Cldn10
Pon1
Rbp4
Elf5
Sftpa1
Scgb3a2
Cxcl15
Mgst3
Tspan1
Gpx2
Epb41l4aos
Snhg12
Ifrd1
Cxcl16
Nr1d1
Dbp
Gipc2
Emp2
Krt8
Foxj1
Mlf1
Nme5
Tmem212
Fam183b
Gm867
Transcription factors
Creb3l1
Esr1
Elf5
Epas1
Sox2
Jun Atf3 Myc
Atf4 Irf1 Relb Trp53 Zkscan5
Klf10 Ddit3 Zbtb2 Dbp Bhlhe41
Rela Nfyc Csrnp1 Tef Zkscan14
Creb5 Rel Maff Nr1d2 Zfp362
Foxo3 Nfkb2 Nr1d1 Irf2 EhfEhf
Klf6 Zfp867 Klf9 Bhlhe40
Mafk Hbp1 Hlf Junb
Foxj1 Barx2 Myb
Zfp474 Phtf1
Cell type
club
club_div
ciliated
DTA⁺_club
DTA⁺_Foxj1⁺
SE
PT
K
Club cell markers
Scgb1c1
Scgb3a2
Ciliated cell markers
Myb
Meig1
Foxj1
Dynlrb2
Expression
Pseudotime

**Fig. S7: Trajectory analysis of lung epithelial cells in the SfSRC mouse model.**

**A.** Diffusion maps of dividing and non-dividing basal and goblet cells with RNA velocity vectors calculated individually for each timepoint. **B.** Diffusion map of dividing and non-dividing goblet and club cells with RNA velocity vectors indicating differentiation of goblet into club cells. **C.** Diffusion map of dividing and non-dividing basal, goblet, and club cells with pseudotime trajectory colored by pseudotime of cells selected to characterize the goblet to club cell trajectory in D. **D.** Smoothed expression heatmap of the top 1,000 altered genes along the differentiation trajectory from goblet to club cells. The names of the top seven genes in each cluster are annotated and transcription factors are marked in orange. All transcription factors of each cluster are listed on the right. PT: pseudotime; SE: scaled expression. **E.** Normalized expression along the goblet to club cell pseudotime trajectory for exemplary genes. The black line is the smoothed expression with the confidence interval shown in gray. **F.** Diffusion map of all secretory cell types. **G.** Diffusion map of dividing and non-dividing club, DTA$^+$_club, DTA$^+$_Foxj1$^+$, and ciliated cells with pseudotime trajectory colored by cell type. **H.** Diffusion map of DTA$^+$_club, DTA$^+$_*Foxj1*$^+$, and ciliated cells with RNA velocity vectors indicating transition of DTA$^+$_club into DTA$^+$_*Foxj1*$^+$ cells that are transitioning into a ciliated-like state. **I.** Diffusion map of dividing and non-dividing club, DTA$^+$_club, DTA$^+$_*Foxj1*$^+$, and ciliated cells colored by pseudotime of cells selected to characterize the club to ciliated cell trajectory in I. **J.** Smoothed expression heatmap of the top 1,000 altered genes along the differentiation trajectory from club to ciliated cells. The names of the top three genes in each cluster are annotated and transcription factors are marked in orange. All transcription factors of each cluster are listed on the right. PT: pseudotime; SE: scaled expression. **K.** Normalized expression along the club to ciliated cell pseudotime trajectory for exemplary genes. The black line is the smoothed expression with the confidence interval shown in gray. Figures were calculated on merged expression data of the SfSRC model from all timepoints (day 0, 2, and 3).

### Supplementary tables provided as excel files

**Supplementary Table 1: Marker genes for dividing cell populations in the lungs of SRC mice after two and three days of tamoxifen expression.** Table shows AUC, power, average logFC, percentage of cells expressing the gene in the population (pct.1) and percentage of cells expressing the gene in the rest of the cells (pct.2). Markers were calculated using the roc test and pct.>0.2.

**Supplementary Table 2: Marker genes for mesenchymal cells in uninjured lungs (day 0) of SRC mice.** Table shows AUC, power, average logFC, pct.1 and pct.2. Markers were calculated using the “roc” test and pct.>0.2.

**Supplementary Table 3: Marker genes for epithelial cells in uninjured lungs (day 0) of SRC mice.** Table shows AUC, power, average logFC, pct.1 and pct.2. Markers were calculated using the “roc” test and pct.>0.2.

**Supplementary Table 4: Differentially expressed genes at day 2, 3, and 4 after tamoxifen administration in each epithelial population of SRC mice.** Table shows p value, average logFC, pct.1, pct.2 and adjusted p value. DEGs were calculated using MAST, average logFC>2 and adjusted p value <0.05.

**Supplementary Table 5: Marker genes for epithelial cells on day 2 of SRC mice.** Table shows AUC, power, average logFC, pct.1 and pct.2. Markers were calculated using the “roc” test and pct.>0.2.

**Supplementary Table 6: Common marker genes in DTA$^+$ cells.** Table shows AUC, power, average logFC, pct.1 and pct.2. Markers were calculated using the “roc” test and pct.>0.2.

**Supplementary Table 7: Differentially expressed genes in DTA$^+$ like cells of COVID-19 patients.** Table shows p value, average logFC, pct.1, pct.2 and adjusted p value. DEGs were calculated using MAST average logFC>1.5 and adjusted p value <0.05.

**Supplementary Table 8: Marker genes for epithelial cells in SfSRC mice (all timepoints merged).** Table shows AUC, power, average logFC, pct.1 and pct.2. Markers were calculated using the “roc” test and pct.>0.2.

**Supplementary Table 9: DEGs during goblet to basal cell differentiation.** Table shows q and Moran’s I values.

**Supplementary Table 10: DEGs during goblet to club cell differentiation.** Table shows q and Moran’s I values.

**Supplementary Table 11: DEGs during club to ciliated cell differentiation.** Table shows q and Moran’s I values.

**Supplementary Table 12: DEGs during club to AT2 cell differentiation.** Table shows q and Moran’s I values.

**Supplementary Table 13: DEGs during AT2 to DTA$^+$ *Sftpc*$^+$ cell differentiation.** Table shows q and Moran’s I values.

**Supplementary Table 14: Ligand-receptor pairs used for crosstalk analysis.** CellChat mouse database ligand-receptor pairs manually curated.

**Supplementary Table 15: Detailed information for all scRNA-seq runs.** Table shows cell type analyzed, mouse model, timepoint, number of cells analyzed, filtering parameters, mean reads per cell, median genes per cell, and median UMI counts per cell.
